# High-Throughput Bioprinting of Spheroids for Scalable Tissue Fabrication

**DOI:** 10.1101/2024.06.30.601432

**Authors:** Myoung Hwan Kim, Yogendra Pratap Singh, Nazmiye Celik, Miji Yeo, Elias Rizk, Daniel J. Hayes, Ibrahim T Ozbolat

**Author notes:** These authors contributed equally to this work (Myoung Hwan Kim, Yogendra Pratap Singh). Author to whom any correspondence should be addressed.

## Abstract

Tissue biofabrication that replicates an organ-specific architecture and function requires physiologically-relevant cell densities. Bioprinting using spheroids has the potential to create constructs with native cell densities, but its application is limited due to the lack of practical, scalable techniques. This study presents HITS-Bio (High-throughput Integrated Tissue Fabrication System for Bioprinting), a novel multiarray spheroid bioprinting technology enabling scalable tissue fabrication by rapidly positioning a number of spheroids simultaneously using a digitally-controlled nozzle array (DCNA) platform. HITS-Bio achieves an unprecedented speed, an order of magnitude faster compared to existing techniques while maintaining high cell viability (>90%). The platform’s ability to pattern multiple spheroids simultaneously enhances fabrication rates proportionally to the size of DCNA used. The utility of HITS-Bio was exemplified in multiple applications, including intraoperative bioprinting with microRNA transfected spheroids for calvarial bone regeneration (∼30 mm^3^) in a rat model achieving a near-complete defect closure (∼91% in 3 weeks and ∼96% in 6 weeks). Additionally, the successful fabrication of scalable cartilage constructs (1 cm^3^) containing ∼600 chondrogenic spheroids highlights its high-throughput efficiency (under 40 min per construct) and potential for repairing volumetric tissue defects.

## 1. Introduction

Three-dimensional (3D) bioprinting has been making a progressive impact on medical sciences, which has great potential to facilitate the fabrication of functional tissues not only for transplantation but also for disease modeling and drug screening. It offers great precision for the spatial placement of cells, which is crucial for guiding tissue repair and regeneration ^1^. Despite the significant progress in bioprinting of cells, the standard approach of encapsulating cells within a hydrogel bioink faces major limitations. One key challenge is achieving physiologically-relevant cell densities (100-500 million cells/mL) ^2^. This limitation hampers the success in development of functional tissues, as the resulting bioprinted constructs lack sufficient cell densities. Indeed, most hydrogel bioinks require a compromise between cell density and polymer density to achieve printability, often resulting in suboptimal conditions for the intended application.

In response to these challenges, tissue spheroids are considered a promising candidate. Spheroids are cellular aggregates that have been utilized in tissue fabrication due to their advantages, including native-like cell density and the capability to secrete substantial levels of extracellular matrix (ECM) components with effective communication among cells in a 3D native-like microenvironment ^3^. Due to these advantages, spheroids have been considered as building blocks for bioprinting a variety of tissues ^3^. Spheroid density and spatial arrangement are critical factors in achieving native-like tissue physiology, as they ensure the precise arrangement and high cell concentration necessary for effective tissue formation ^4^. Despite the progress in spheroid bioprinting technologies, the major shortcomings associated with them, such as poor positioning of spheroids, significant loss of viability and structural integrity, poor repeatability of the process when using non-uniform size spheroids, inability to form complex 3D shapes, and most importantly, the lack of scalability, limit their translation. Contemporary techniques, such as extrusion-based bioprinting ^5^, droplet-based bioprinting ^6^, Kenzan ^7^, and bio-gripper approaches ^8^, have been utilized to bioprint spheroids as building blocks; however, most of them suffer from poor positioning of spheroids, significant loss of their viability and structural integrity, poor process repeatability, and the lack of scalability. Previously, a high-precision technique, known as ‘aspiration-assisted bioprinting (AAB)’ ^9^ was introduced, where, an aspiration force is applied to position a biologic (e.g., cells, tissue spheroids or strands) during the bioprinting process. This technique enables picking and placement of biologics ranging from 80 to 800 μm in size into or onto a gel substrate with minimal cellular damage (>90% cell viability) and achieving high positional precision (∼11% with respect to spheroid size). AAB has recently inspired applicability in magnetic lifting of neural organoids for the construction of assembloids ^10^, 3D printing of living moving organisms (i.e., beetles ^11^) and bioprinting of high cell-density tissues (i.e., cartilage ^12^ and bone ^13^) and disease models (post-myocardial infarction scarring) ^14^. However, the major limitation of this technique is its reliance on bioprinting one spheroid at a time. While this one-at-a-time approach allows for multiple iterations in the deposition of spheroids to scale up, it significantly prolongs the bioprinting process (∼20 sec/spheroid), similar to other spheroid bioprinting techniques ^15^, which poses major challenges to increasing the scalability of fabricated tissues consistently and efficiently.

In this work, we address a long-standing problem in 3D bioprinting of spheroids and demonstrate a novel technology named “HITS-Bio: High-throughput Integrated Tissue fabrication System for Bioprinting”. HITS-Bio represents a significant advancement in rapid bioprinting of spheroids for scalable tissue fabrication. This technique enabled the bioprinting of scalable tissues via precise positioning of spheroids (and also applicable to organoids) in a high-throughput manner at an unprecedented speed (an order of magnitude faster than the existing techniques) with high cell viability (>90%) using a digitally controlled nozzle array (DCNA) for the patterning and spatial distribution of several spheroids simultaneously. The capacity of the DCNA platform to pattern several spheroids at a time enabled the rapid creation of tissues, thus increasing the fabrication rate by a factor of *n* (the number of nozzles in DCNA) proportionally to the number of nozzles used. Herein, we presented an innovative bioink to support spheroids during bioprinting, analogous to assembling building blocks where the bioink acts as cement and spheroids serve as bricks. To demonstrate the practical application of this technology, we first focused on calvarial bone regeneration, where critical size defects were repaired through intraoperative bioprinting (IOB) of osteogenically-committed bone spheroids. The study introduced intraoperative bioprinting with spheroids for the first time, enabling on-demand tissue fabrication and reducing surgery time. It innovatively used micro-RNA (miR) technology to achieve osteogenic differentiation of spheroids, and HITS-Bio enabled simultaneous or sequential aspiration and bioprinting of miR-transfected spheroids on demand. Moreover, the potential of HITS-Bio in the context of scalable tissue fabrication was exemplified through the successful creation of cm^3^ cartilage tissue constructs, which were precisely assembled using ∼600 chondrogenically committed spheroids under 40 min per construct, representing a scale of fabrication with a high efficiency that surpasses the capabilities of current bioprinting technologies.

## 2. Results

### 2.1 Working mechanism of the HITS-Bio process

The HITS-Bio platform featured a facile assembly inside a biosafety hood due to its compact footprint, designed as a table-top system equipped with various accessories such as cameras for detailed observation and analysis. The platform had three main components: an innovative multinozzle DCNA (**Figure S1**), a high-precision XYZ linear stage to move DCNA in 3 axes (X, Y, and Z), and an extrusion head to deposit a gel substrate (**Figure S2**). It was operated by a custom-made software interface (**Supplementary Video 1)**. To visualize the bioprinting process in real-time and verify the actual position of DNCA in 3D, three microscopic cameras for the isometric, bottom, and side views were integrated (**Figure S2**). DCNA facilitated picking up multiple spheroids by controlling the aspiration pressure in the selectively opened nozzle depending on the design of applications, as demonstrated in **Figures 1A** and **S3**. To lift spheroids, DCNA was moved to a Petri dish, where spheroids were suspended in a culture medium. After spheroids were successfully aspirated to the end of the selectively opened DCNA, confirmed by the bottom view camera, DCNA with spheroids was gently lifted from the spheroid chamber (**Figure 1A**, Step 1.4). As a substrate to place the spheroids, a bioink was extruded (**Figure 1B**, Step 2.1). Next, DCNA loaded with spheroids was transferred over the substrate (**Figure 1B**, Steps 2.2 and 2.3). Once the spheroids were in contact with the substrate, aspiration pressure was cut off to deposit the spheroids (**Figure 1B**, Step 2.4). We exemplified the utility of HITS-Bio in three different configurations. For in-vitro bioprinting of single-layer-spheroids, low density (composed of 16 spheroids) and high density (composed of 64 spheroids) were bioprinted by repeating the process as depicted in **Figures 1A-B** (Steps 1.1 to 1.4 and from Steps 2.2 to 2.4). After the spheroid placement, another layer of bioink was deposited on top of the bioprinted spheroids to envelop them, followed by photo-crosslinking using a 405 nm LED light source for 1 min (**Figure 1B**, Step 2.5). For IOB (**Figure 1C**), after creating a critical-sized calvarial defect, DCNA was positioned over it. The bone ink (BONink) was extruded at the defect area and spheroids were deposited at two different spheroid densities (low - 16 spheroids and high - 64 spheroids). Then, another layer of BONink was extruded over the spheroids (**Figure 1C**, Step 3.6), followed by photo-crosslinking and suturing of the skin. For scalable tissue bioprinting (**Figure 1D**, Steps 4.1 – 4.5), scalable cartilage tissues (SCTs) were created, where a cartilage ink (CARink) was extruded followed by the precise placement of 64 chondrogenic spheroids. This iterative process was repeated nine times to assemble a construct comprising 9 stacked tissue layers and a total of 576 spheroids.

**Figure 1.**
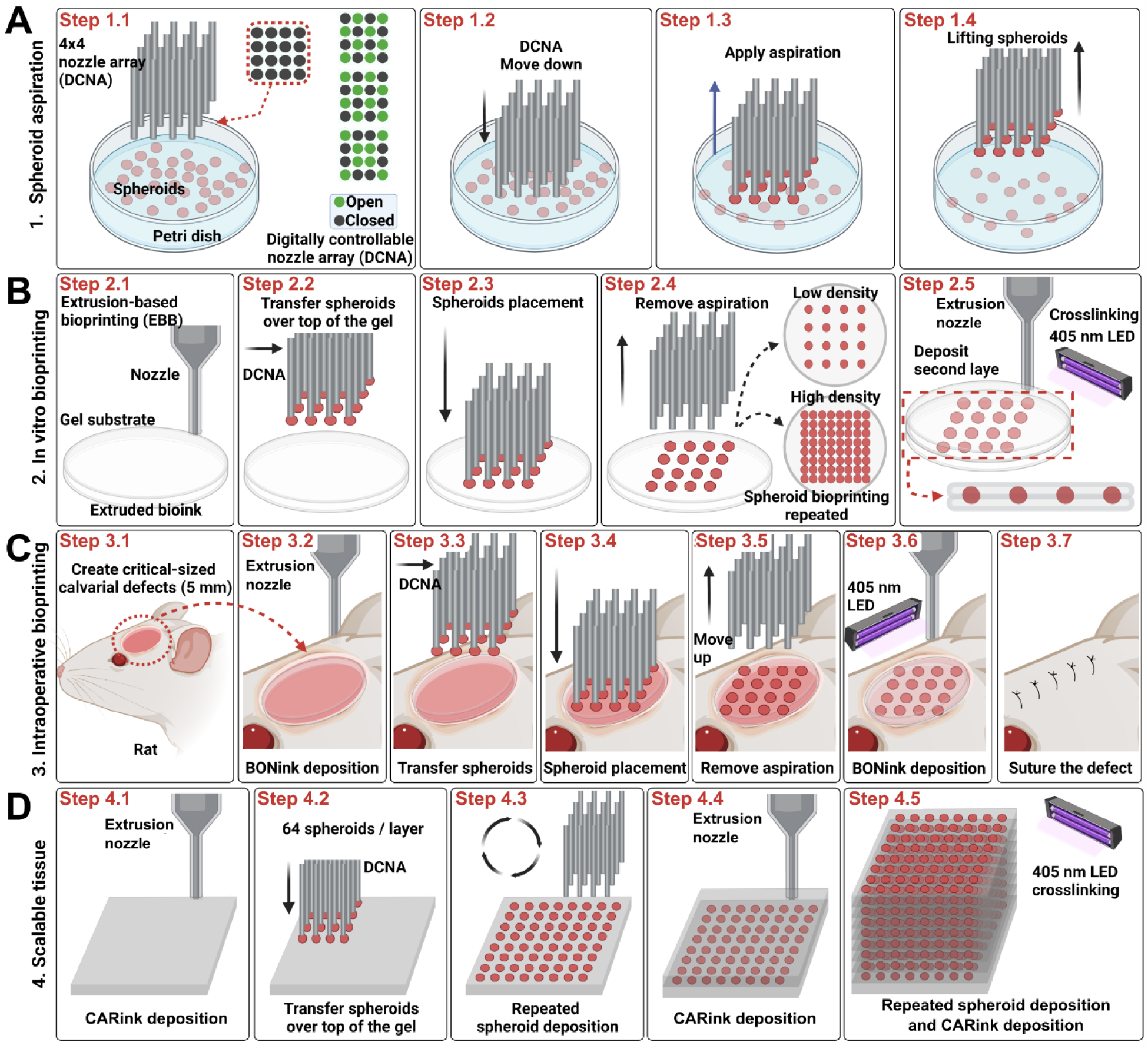
Schematic of the HITS-Bio process. (A) Spheroid loading with DCNA, which selectively enabled or disabled aspiration according to a user-defined design in an iterative manner. (B) *In vitro* bioprinting of spheroids including extruded bioink via EBB and spheroid placement using DCNA, having different spheroid loading densities (i.e. low density – 16 spheroids and high density – 64 spheroids) in an iterative manner. (C) IOB of spheroids into a rat calvarial defect, where spheroids were sandwiched between the extruded BONink layers. (D) 1 cm^3^ of SCT bioprinted with CARink and ∼600 spheroids. Created with BioRender.com.

DCNA consisted of stainless-steel needles with predefined spaces between them, which were assembled by precisely stacked multiple acrylic plates (**Figure S1**). A micro-manufactured 4x4 nozzle array with different configurations (various designs with widths from 2.8 to 4.0 mm) was prepared as shown in **Figure S4A**. To calculate the manufacture inaccuracies during micro-manufacturing of DCNA, bottom and side views of DCNA were captured. These views allowed the measurement of inter-nozzle distance and accuracy of each nozzle against a reference point. The range of widths was determined based on the area, where spheroids could be bioprinted in a circular area with a diameter of 5 mm. As shown in **Figures 2A** and **2B**, the actual width and inter-nozzle distance of DCNA were measured and compared with the designed width and inter-nozzle distance (black dots in **Figures 2A** and **2B**), respectively. Considering the error expected during laser cutting, DCNA was micro-manufactured with less than 5% error. The positional XY (**Figure S4C**) and Z (**Figure 2C**) errors of DCNA were less than 5% for 300 µm spheroids tested in this study. We observed that lifting spheroids from the culture medium into air resulted in entrapment of liquid (culture medium) between nozzles and its elevation from its surface acted upon by capillary forces due to its surface tension between the liquid and DCNA (**Figure S4B, Supplementary Video 2**). The elevated liquid hindered successful spheroid lifting and placement. For example, comparing two different sizes of DCNA (2.8 v.s. 3.4 mm), higher liquid elevation was observed in the closely packed 2.8-mm DCNA, where spheroids experienced resistance in their lifting due to the elevated liquid. In contrast, the 3.4-mm DCNA showed a much lower height of liquid elevation, which resulted in successful spheroid lifting (**Supplementary Video 2**). Thus, optimizing parameters, such as inter-nozzle distance and liquid elevation, was crucial in relation to the likelihood of successful spheroid lifting.

**Figure 2.**
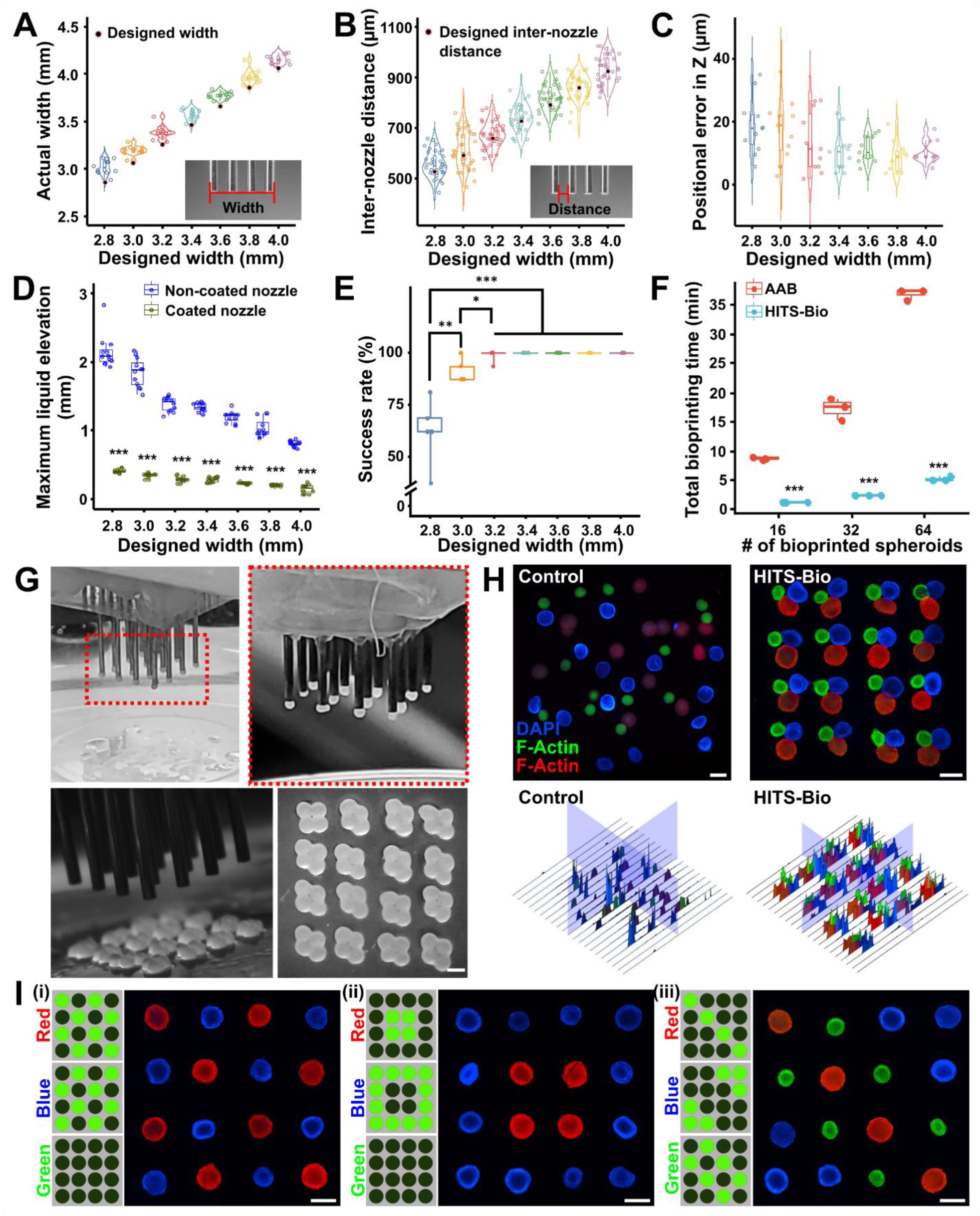
Analysis of HITS-Bio performance. Measurement of (A) end-to-end nozzle array width (*n* = 12), (B) inter-nozzle distance (*n* = 36), and (C) positional error in the Z-axis (*n* = 11). (D) Maximum liquid elevation in DCNA (*n* = 12, \*\*\**p* < 0.001). (E) The impact of interspatial distances of silicon-coated DCNA on the success rate of spheroid lifting (*n* = 5, \**p* < 0.05, \*\**p* < 0.01, and \*\*\**p* < 0.001). (F) A comparison of elapsed bioprinting time between conventional single-nozzle aspiration-assisted bioprinting (AAB) and HITS-Bio (*n* = 3, \*\*\**p* < 0.001). (G) Lifted spheroids from a spheroid chamber and placed on a GM10 substrate in an iterative manner to bioprint 64 spheroids to generate a rhombus pattern. Scale bar: 300 μm. (H) Fluorescence images and the corresponding intensity map of three different sizes and colors (stained with DAPI (Blue), F-Actin (Red), F-Actin (Green)) of spheroids (16 spheroids per color) using manually mixed (control) and HITS-Bio. Scale bar: 500 μm. (I) Selectively patterned spheroids stained with DAPI (blue), F-Actin (red), and F-Actin (green) using the DCNA platform with various configurations. Scale bar: 500 μm.

During lifting, a capillary reaction was observed between nozzles, where liquid rose in a narrow gap against gravity (**Figure S4D**). At equilibrium, the upward force due to surface tension balances the downward force of the liquid weight. The liquid continues to rise until these forces are equal. As detailed in **Supplementary Information S1**, this relationship can be expressed mathematically, displaying that the height of the elevated liquid is inversely proportional to the inter-nozzle distance and surface tension. To reduce the height of the elevated liquid, one can either increase the inter-nozzle distance or decrease the surface tension. Subsequently, various inter-nozzle distances of DCNA were explored, showing correlation between the quantified elevated liquid length with different sizes of DCNA. Subsequently, silicon coating was applied to DCNA to lower the surface tension by reducing the surface energy. The results indicated that the liquid elevation was significantly decreased after the silicon coating on DCNA surface (**Figure 2D**). In addition, the silicon-coated DCNA was lifted smoothly from the liquid, as opposed to the non-coated DCNA, which tended to drag the liquid (**Supplementary Video 2**). Furthermore, as the inter-nozzle distance decreased, the liquid elevation increased. Thus, the success rate of spheroid lifting decreased with reducing designed width due to the liquid elevation (**Figure 2E**). In other words, a silicon-coated DCNA with a 3.4-mm designed width showed complete success of spheroid lifting (**Figure 2E and Supplementary Video 3**) and was selected for further consideration in the HITS-Bio process.

To test the performance of HITS-Bio with the optimized DCNA, we benchmarked it against a single nozzle AAB system (**Figure 2F**). The results indicated that HITS-Bio was highly efficient when a large number of spheroids were needed to be bioprinted. Using DCNA with 16 nozzles (**Figure 2G**), the bioprinting process for 16 spheroids took less than 1 min, which was significantly faster than the conventional AAB system, which took nearly an order of magnitude longer (**Figure 2F**) despite the AAB system being upgraded for its bioprinting speed (with full automation) compared to previous publications ^9,12^. As the number of bioprinted spheroids increased, the disparity in total bioprinting time between AAB and HITS-Bio increased exponentially (**Figure 2F and Supplementary Video 4**). To evaluate the performance of HITS-Bio for the patterning of spheroids, various types of spheroids were selectively lifted and patterned on a gelatin methacryloyl (GM) substrate. For example, we compared HITS-Bio with manual loading of spheroids (**Figure 2H**). In this regard, a total of 48 spheroids (16 Green (∼350 μm in diameter), 16 Blue (∼425 μm in diameter) and 16 Red (∼500 μm in diameter)) per sample were utilized. To prepare the control group, spheroids were mixed in 10% GM using a pipette in a 3D-printed mold (**Figures S5A-B**). For HITS-Bio, spheroids were bioprinted into the GM substrate in a triplet arrangement, where triplets were patterned with precise control on their positioning while the manually-loaded spheroids resulted in random distribution with a lack of control, not just in X- and Y-axis but also Z-axis as most of the spheroids were not in the focal plane (**Figure 2H**). HITS-Bio showed 100% spheroid loading efficiency, meaning all spheroids were bioprinted and loaded into GM successfully regardless of their size. In contrast, 80-85% efficiency was attained in the case of manual loading (**Figures S5C and S5D**). The reduced efficiency in manual loading was due to the fact that spheroids were sticking to the wall of the pipette tip during manual deposition (**Figure S5B**). One would expect that the efficiency would further decrease as the number of spheroids increased. Therefore, using HITS-Bio, the number and type of spheroids could be controlled to precisely position them with 100% loading efficiency. To further confirm the spatial placement of spheroids using HITS-Bio, the three types of spheroids were patterned in various configurations (**Figure 2I**). Herein, based on the desired location of spheroids, the corresponding valves in DCNA were digitally activated and switched on, and the spheroids were loaded on the nozzles selectively as shown in **Supplementary Video 5**. The spheroids were released in the designated positions on demand. Therefore, HITS-Bio facilitated the successful patterning of different types of spheroids at desired positions regardless of their size without loss of spheroids.

### 2.2 *In vitro* development and characterization of bioinks as a substrate for spheroid bioprinting

To develop innovative bioinks akin to versatile "cement-like substrate" capable of assembling spheroids (bricks) into structured building blocks for bioprinting, components were chosen for their compatibility with HITS-Bio in terms of their extrusion, the ability to form a slightly adhesive cement-like substrate to retain spheroids in desired patterns, and to enhance ECM formation. The bioink base was composed of GM for its biocompatibility and tunable mechanical properties, nanohydroxyapatite (HA) to enhance osteoconductivity, and β-glycerophosphate disodium salt hydrate (β-GP) for thermal gelation. Hyaluronic acid sodium salt (HyA) was added to improve cell adhesion and proliferation, fibrinogen for its role in promoting cell-matrix interactions, and lithium phenyl (2,4,6-trimethylbenzoyl) phosphinate (LAP) as a photoinitiator to enable light-induced crosslinking, ensuring structural integrity and stability of the bioink. These components were mixed in various ratios to obtain composite bioinks, where the bone ink (**Figure S6A**) was formulated with the same components as the cartilage ink (**Figure S6B**), but with the addition of β-GP and HA. GM at a concentration of 10% was denoted as GM10, and at 20% it was labeled GM20. Similarly, HA at 15% concentration was designated as HA15, and at 30%, it was labeled HA30. To assess the bioinks, their rheological properties, mechanical behavior, and degradation profile were evaluated. Rheological analysis was performed to test the flow characteristics of the bioinks. The viscosity profile, determined through a flow sweep across varying shear rates from 0.1 to 100 s^−1^, revealed a shear thinning behavior in all bioink formulations, making them suitable for extrusion (**Figure 3Ai**). Bioinks with higher concentrations of GM (GM20) and HA (HA30) exhibited higher viscosities compared to GM10 and HA15 formulations, respectively. The recovery sweep test (**Figure 3Aii**) showed that the viscosity of all samples recovered rapidly, suggesting that the bioinks exhibited a self-healing property. The viscosity profile of the bioinks during the recovery followed the same trend as that of the flow sweep. Further, the frequency sweep showed that storage modulus (G′) was higher than loss modulus (G″) in the frequency range of 0.1–100 rad/s (**Figure 3Aiii**). G′ and G″ were independent of frequency for the GM-alone samples, while they were slightly dependent for the composite bioinks. The compression testing of bioinks exhibited a non-linear stress-strain curve. Among the GM20 samples, GM20HA30 exhibited the highest modulus at 360.7 ± 54.3 kPa, followed by GM20HA15 at 221 ± 27.7 kPa, and GM20 at 154.7 ± 18.4 kPa. Conversely, GM10 samples displayed lower moduli, with values of 24 ± 3.3 kPa, 22.6 ± 4.2 kPa, and 3.8 ± 1.7 kPa for GM10HA30, GM10HA15, and GM10, respectively (**Figure 3B**). Although GM20 samples exhibited a higher modulus, they demonstrated a lower fracture strain ranging from 40-45%, whereas the GM10 samples fractured at a higher strain of 65-70% (**Figure S6C**).

**Figure 3.**
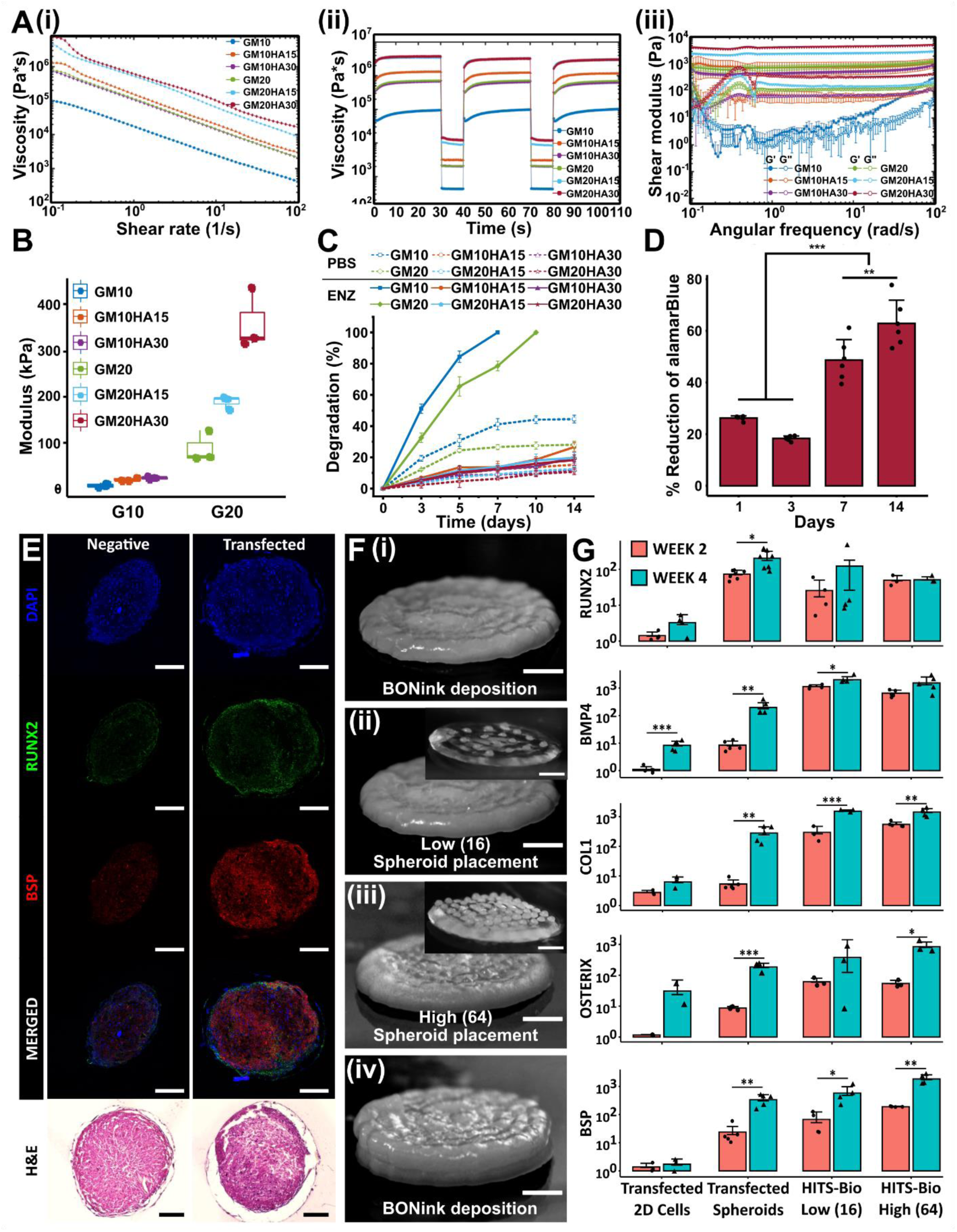
In vitro characterization of bioinks. (A) Rheological characterization of BONink and CARink at different concentrations of HA and GM (*n* = 3): (i) Flow curves of bioinks from the rotational test at a shear rate ranging from 0.1 to 100 s^-1^ showing mean plots. (ii) Recovery behavior and viscosity measurements at five intervals at alternating shear rates (shear rate of 0.1 s^-1^ for 30 s and 100 s^-1^ for 10 s) showing mean plots. (iii) Measured shear modulus from frequency sweeps at an angular frequency ranging from 0.1 to 100 rad s^-1^; with error bars showing mean ± s.d. (B) Compressive modulus (*n* = 3), (C) % degradation of samples for 14 days in enzyme (ENZ) and PBS (*n* = 3), and (D) % reduction of alamarBlue to assess cell viability (*n* = 6, \*\**p* < 0.01 and \*\*\**p* < 0.001) with G20HA30 for 14 days. (E) Histomorphometric characterization of non-transfected and miR-(196a-5p + 21) co-transfected spheroids stained with RUNX2, BSP, and H&E at Week 4. (F) Steps involved during the bioprinting process for *in vitro* fabrication of bone constructs at two different spheroid densities: (i) BONink deposition, (ii) bioprinted spheroids at low (16 spheroids) and (iii) high density (64 spheroids), and (iv) covered the bioprinted spheroids with another layer of BONink. Scale bar: 1 mm. Inset images demonstrate bioprinted spheroids with low and high densities on a transparent gel (CARink) for clear visualization. Scale bar: 1 mm. Scale bar: 200 µm. (G) Quantification of RUNX2 (*n* = 4, 2D transfected; *n* = 7, transfected spheroids; *n* = 4, low (16 spheroids); and *n* = 3, high (64 spheroids)), BMP-4 (*n* = 5, 2D transfected; *n* = 5, transfected spheroids; *n* = 4, low (16 spheroids); and *n* = 5, high (64 spheroids)), COL-1 (*n* = 3, 2D transfected; *n* = 5, transfected spheroids; *n* = 3, low (16 spheroids); and *n* = 5, high (64 spheroids)), OSTERIX (*n* = 2, 2D transfected; *n* = 4, transfected spheroids; *n* = 3, low (16 spheroids); and *n* = 3, high (64 spheroids)), and BSP (*n* = 3, 2D transfected; *n* = 5, transfected spheroids; *n* = 4, low (16 spheroids); and *n* = 4, high (64 spheroids)) gene expression of bioprinted bone (\**p* < 0.05, \*\**p* < 0.01 and \*\*\**p* < 0.001).

The degradation test results indicated a decrease in enzymatic degradation rate for GM samples upon HA incorporation, with complete degradation occurring in ∼7 days for 10% GM and 10 days for 20% GM (**Figure 3C**). The HA-based composites demonstrated limited degradation, with only ∼18% for GM20HA30 and ∼20% for GM20HA15 over 14 days. Similarly, in GM10 samples, degradation amounted to ∼20% for GM10HA30 and ∼27% for GM10HA15. Conversely, composite bioinks in phosphate-buffered saline (PBS) only underwent 12-15% degradation, while only GM samples in the same medium showed higher degradation levels of 30-40%. After characterizing bioinks for their rheological, mechanical, and degradation properties, the GM20 and GM20HA30 composite were selected and named CARink (cartilage ink) and BONink (bone ink), respectively, for further investigations. GM10 and its composites exhibited poor printability and mechanical properties, whereas GM20HA15 demonstrated inadequate mechanical strength. After developing the BONink, the metabolic activity of hADSCs in the BONink was assessed over time, which revealed significantly higher proliferation observed on Day 14 compared to Day 7 (*p* ≤ 0.01), and lower levels on Days 1 and 3 (*p* ≤ 0.001) (**Figure 3D**). These results demonstrate the biocompatibility of the BONink, affirming its suitability for bone tissue engineering applications. Osteogenically-committed spheroids were then formed using miR-transfected hADSCs as shown in **Figure S7A**. Specifically, hADSCs were transfected with miR-196a-5p, or miR-21, or in combination (miR-(196a-5p + 21)). Transfection was validated via gene expression (**Figure S7B**) and histomorphometric characterization (IHC and H&E) (**Figures 3E and S8**) of osteogenic markers at Weeks 2 and 4. In our preliminary experiments, transfection was assessed in 2D conditions, where hADSCs with and without the miR-transfection were evaluated, and the differentiation in established osteogenic medium was taken as positive control. The results of preliminary tests showed upregulated osteogenic genes (*RUNX2*, *BMP-4*, *COL-1*, *OSTERIX*, and *BSP*) when hADSCs were co-transfected with miR-(196a-5p + 21) compared to miR-196a-5p alone and non-transfected hADSCs (negative control) (**Figure S7B**). The expressions were significantly higher (*p* ≤ 0.05) at Week 4 as compared to Week 2 and comparable to the positive control. Additionally, IHC assessment of RUNX2 and BSP markers showed intense staining in the miR-(196a-5p + 21) transfected hADSCs, which was comparable to the positive control and higher than the miR-196a-5p alone and non-transfected hADSCs (**Figure 3E and S8**). These results were in agreement with qPCR results. Further, H&E staining showed a strong dark purple color indicating bone mineralization in the positive group (**Figure S8**). The transfected group also showed darker color compared to the negative control and miR-196a-5p alone transfected hADSCs. Overall, miR-transfection supported the differentiation of hADSCs into an osteogenic lineage. Based on these results, for the rest of this study, the co-transfection of miR-(196a-5p + 21) was used for osteogenic differentiation of hADSCs for use in spheroids as building blocks for bone tissue fabrication.

Figure 3F demonstrates the steps for the fabrication of single-spheroid-layer bone constructs with two different densities (16 or 64) of spheroids. Spheroids (350 µm) were placed with a targeted inter-spheroid distance of 670 and 160 µm for the low density (16) and high density (64) spheroids, respectively. The BONink was first extruded in a spiral pattern with 100% infill to lay down a gel substrate with a diameter of ∼ 5 mm prior to spheroid placement (Figure 3Fi). On the BONink, spheroids were bioprinted at low (Figure 3Fii) and high (Figure 3Fiii) densities. After the spheroid placement, another layer of BONink was extruded to overlay spheroids (Figure 3Fiv). It is pertinent to note that the BONink deposition did not displace the previously placed spheroids considerably (**Supplementary Videos 6 and 7**). The final constructs were photo-crosslinked with a 405 nm light source for 1 min and incubated thereafter. qPCR results indicated increased expression of osteogenic markers in the transfected non-bioprinted and bioprinted (low or high density) spheroids (Figure 3G). There was a significant increase across all groups at Week 4 compared to Week 2. As shown in **Table S1**, the results revealed that *RUNX2*, an early osteogenic marker, increased up to 4.8-fold, with the highest levels observed in the low-density group. Similarly, *BMP-4*, another early marker, showed a maximum increase of 23.3-fold in transfected spheroids. *COL1*, involved in both early and late stages of osteogenic differentiation, increased by up to 52.1-fold in transfected spheroids. *OSTERIX*, an intermediate to late-stage marker, increased up to 15.4-fold in the high-density spheroids group. Finally, *BSP*, a late-stage marker crucial for bone strength, increased by 14.1-fold in the transfected spheroids at Week 4 compared to Week 2. These results indicate that the bioprinted constructs, particularly those with lower spheroid density, show increased early osteogenic activity (*RUNX2* and *BMP-4*). In contrast, high-density spheroids and transfected spheroids exhibit significant increases in both intermediate (*OSTERIX*) and late-stage (*BSP*) markers, suggesting more mature osteoblast activity and active-matrix mineralization. Furthermore, the gene expression results were corroborated by the IHC staining, where RUNX2 and OSTERIX markers exhibited expression profiles consistent with their roles in osteoblast differentiation (**Figure S9**). RUNX2 displayed a distinct increase in staining in the low-density group at Week 4 while the high-density group maintained its initial high intensity. On the other hand, OSTERIX showed intense staining in bioprinted constructs particularly in the high-density group, indicating successful active osteoblast differentiation. The IHC staining patterns aligned well with the observed fold-changes in gene expression, reinforcing the dynamic nature of osteogenic differentiation and the sequential involvement of related genes.

### 2.3 Intraoperative bioprinting for bone regeneration in calvarial defects

Following the *in vitro* assessment of physical properties, printability, cytocompatibility, and osteogenic potential of the BONink, we performed animal studies using a rat model. Minimizing the animal numbers, two critical-sized calvaria defects, each 5 mm in diameter, were created on either side of the parietal bone on the rat skull and IOB was performed under aseptic surgical settings (Figure 4A **and Figure S2A**). Akin to *in vitro* settings, the BONink was extruded at the defect site in a spiral pattern with 100% infill (**Supplementary Video 8**). Four groups were considered including (i) empty defect (control), (ii) BONInk only, (iii) low density (16 spheroids), and (iv) high density (64 spheroids). Post IOB, bone regeneration was evaluated at 3 and 6 weeks. Micro-computed tomography (µCT) results (Figure 4B) revealed that the high-density group consisting of 64 spheroids exhibited superior bone regeneration, closing almost the entire defect in Week 6. In contrast, bone regeneration in the empty control group was mainly confined to the periphery of the defects. As a quantitative metric to assess the efficacy and extent of bone tissue formation, bone volume to total volume (BV/TV) was calculated, which revealed significantly higher bone regeneration of ∼38 and 33% in Week 3, and ∼39 and 39% in Week 6 for low- and high-density group, respectively, compared to the BONink-only (*p* ≤ 0.05) and empty group (*p* ≤ 0.01) (Figure 4Ci). Moreover, the normalized bone mineral density (BMD), reflecting the density of regenerated bone normalized to the native bone density, exhibited ∼29 and 28% at Week 3, and ∼34 and 34% at Week 6 for low and high-density groups, respectively, compared to the BONink-only (p ≤ 0.05) and empty group (*p* ≤ 0.01) (Figure 4Cii). Additionally, the bone coverage area (%) for the low-density and high-density groups was ∼90 and 91% at Week 3, and ∼88 and 96% at Week 6, respectively (Figure 4Ciii). Moreover, the maximum intensity projections generated from µCT data (Figure 4B) of each group were used for the scoring (1-4) of bony bridging across the defect. The results showed significantly higher scores for spheroid-involved groups compared to empty defect and BONink-only groups at both Weeks 3 and 6 (Figure 4Civ). Furthermore, given the significant role of mechanical properties in bone strength, the regenerated bone samples were subjected to a push-out test following their retrieval after Week 6 (Figure 4D). The high-density group exhibited a significantly higher shear yield strength, ∼1.5 times greater than other groups (*p* ≤ 0.05), indicating enhanced resistance to plastic deformation or failure. Similarly, the modulus of resilience in the high-density group was significantly higher, ∼2.4 times greater than the other groups (*p* ≤ 0.01), signifying higher energy absorption capacity up to the point of yielding. However, there was no significant difference observed among the groups in the shear modulus, representing the material’s resistance to shear deformation.

**Figure 4.**
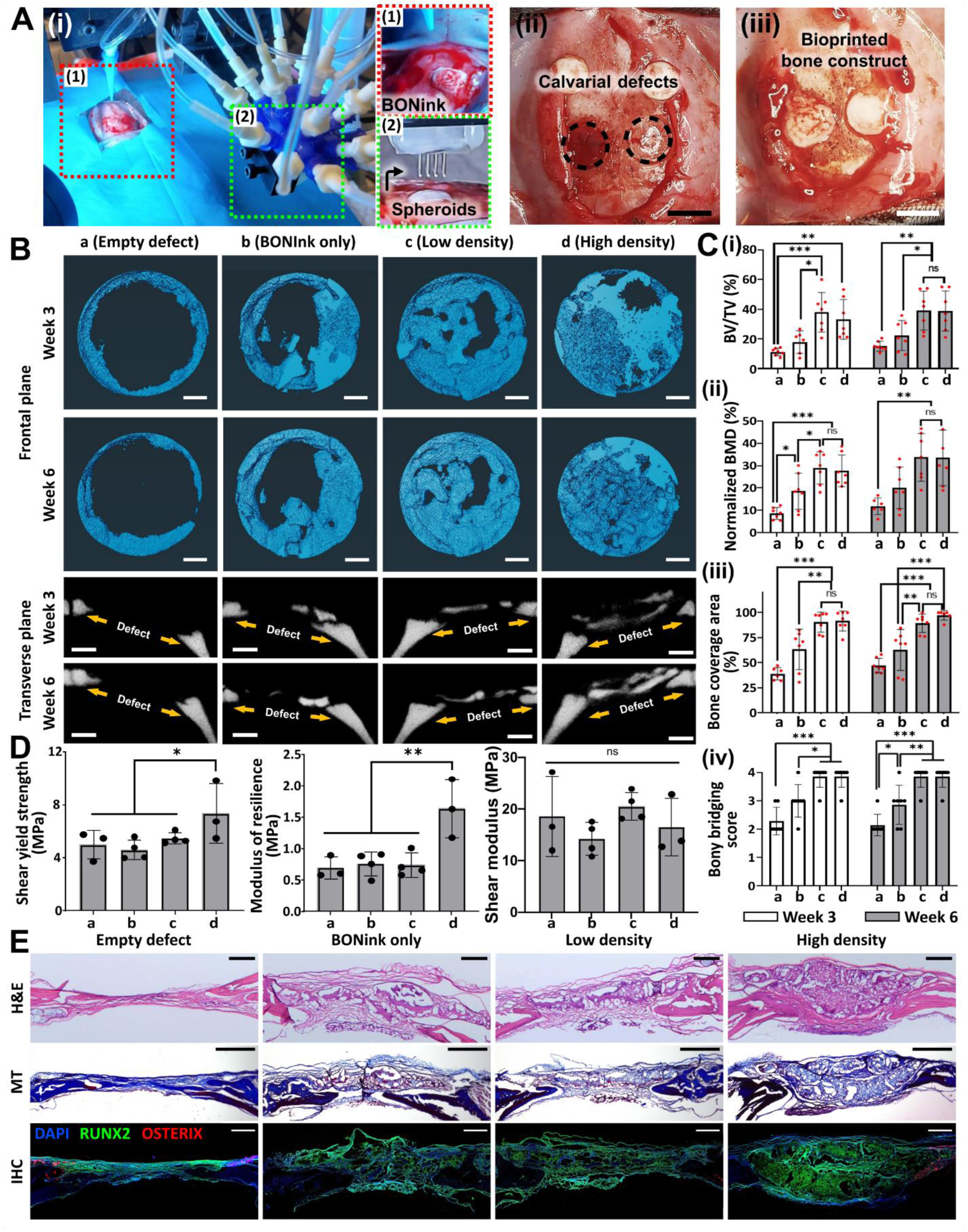
IOB of bone constructs into critical-sized rat calvarial defects for bone regeneration. (A) HITS-Bio setup under surgical settings. Inset images (1-2) demonstrate deposited BONink and spheroid placement, (ii) created calvarial defects (∼ 5 mm in a diameter), and (iii) bioprinted bone constructs with BONink and spheroids. Scale bar: 5 mm. (B) Visualization of newly regenerated bone in calvarial defects with frontal and transverse planes at Weeks 3 and 6 via µCT, including empty, BONink only, low-density and high-density groups. Scale bar: 1 mm. (C) Relevant quantification for bone regeneration, including (i) BV/TV (new bone volume to total bone volume, %), (ii) normalized BMD (bone mineral density, %), (iii) bone coverage area (%), and (iv) scores for bony bridging within the defect examined at Weeks 3 and 6 (*n* = 7, \**p* < 0.05, \*\**p* < 0.01, and \*\*\**p* < 0.001). (D) Mechanical properties of the retrieved defect area 6 weeks after the surgery (Min *n* = 3; \**p* < 0.05, \*\**p* < 0.01 and *ns* not significant). (E) Histomorphometric characterization of sectioned defect after decalcification and stained for H&E (scale bar: 500 µm), MT (scale bar: 1 mm), and IHC (RUNX2 and OSTERIX) (scale bar: 500 µm).

Decalcified sections were then stained with H&E and MT to evaluate the morphology of the regenerated bone (Figure 4E). The H&E images of the high-density group exhibited bridging of the calvarial defect and thicker regenerated bone compared to the soft tissue observed in other groups. MT staining predominantly showed soft tissue formation in the empty defect group, while both spheroid-containing groups showed signs of immature bone formation, which was increased in the high-density group. Thus, histological evaluations of the defects showed that high-density samples illustrated bone formation in 6 weeks, along with active bone healing and mineral deposition. The repair site exhibited characteristics like intramembranous bone formation, aligning with the fact that the development of calvaria is primarily associated with intramembranous ossification ^16^. Furthermore, IHC staining indicated higher expression levels of RUNX2 and OSTERIX in the bioprinted groups compared to the empty defect and BONink-only groups (Figure 4E). Overall, the *in vivo* results showed that the high-density group significantly enhances bone regeneration in critical-size calvarial defects, achieving nearly full defect closure in 6 weeks and superior new bone formation and strength.

### 2.4 Fabrication of scalable cartilage tissues (SCTs)

To illustrate the potential of HITS-Bio in generating volumetric tissues, we fabricated a SCTs (as shown in Figure 1D and **Supplementary Video 9**). For this purpose, the CARink (i.e. GM20 ink) was used as a bioink along with miR-(140 + 21) co-transfected chondrogenic spheroids (**Figure S10A**), as this combination of miRs has shown promising results for cartilage regeneration in a previous study ^17^. To fabricate the constructs, we extruded a layer of CARink, followed by the precise placement of 64 chondrogenic spheroids. This iterative process was repeated nine times to assemble a construct with a volume of 1 cm^3^, comprising 9 stacked tissue layers and a total of 576 spheroids taking under 40 min (Figure 5A). SCTs were then assessed for cell viability and cartilaginous ECM formation. LIVE/DEAD staining revealed that the transfection and tissue fabrication did not impair the cell viability in spheroids, with bioprinted transfected spheroids exhibiting over 90% viability, comparable to non-transfected spheroids (Figures 5B and **5E**). Further, SCTs with co-transfected spheroids exhibited a significantly higher sulfated glycosaminoglycan (sGAG) content compared to the non-transfected control group (*p* ≤ 0.001) at Week 2 (Figure 5C). Interestingly, the DNA content was significantly lower (*p* ≤ 0.001) in the transfected samples compared to the non-transfected samples (Figure 5D). At Week 2, the identification and morphology of chondrogenic spheroids within bioprinted constructs were assessed using H&E. The staining showed the presence of spheroids in the CARink bioprinted constructs, with the chondrogenic spheroids exhibiting more intense staining, indicative of a higher density of ECM deposition and displayed a characteristic cobblestone-like morphology in transfected group at Week 2 (Figure 5F). Additionally, qualitative evaluation of proteoglycans and sGAG was conducted using toluidine blue (TB) staining. In agreement with the quantified sGAG expression (Figure 5C), images of transfected spheroids stained with TB exhibited intense staining, indicating a higher level of sGAG deposition compared to non-transfected samples (Figure 5G). From H&E and TB staining, the co-transfected group indicated chondrogenic lacunae-like properties. IHC staining was also performed to identify the expression of chondrogenic markers, ACAN and COLII, in SCTs. The findings revealed strong fluorescence intensity indicating significantly elevated levels of ACAN (*p* ≤ 0.01) and COLII (*p* ≤ 0.001) in SCTs at Week 2, suggesting the formation of new chondrogenic ECM (**Figure S10B**).

**Figure 5.**
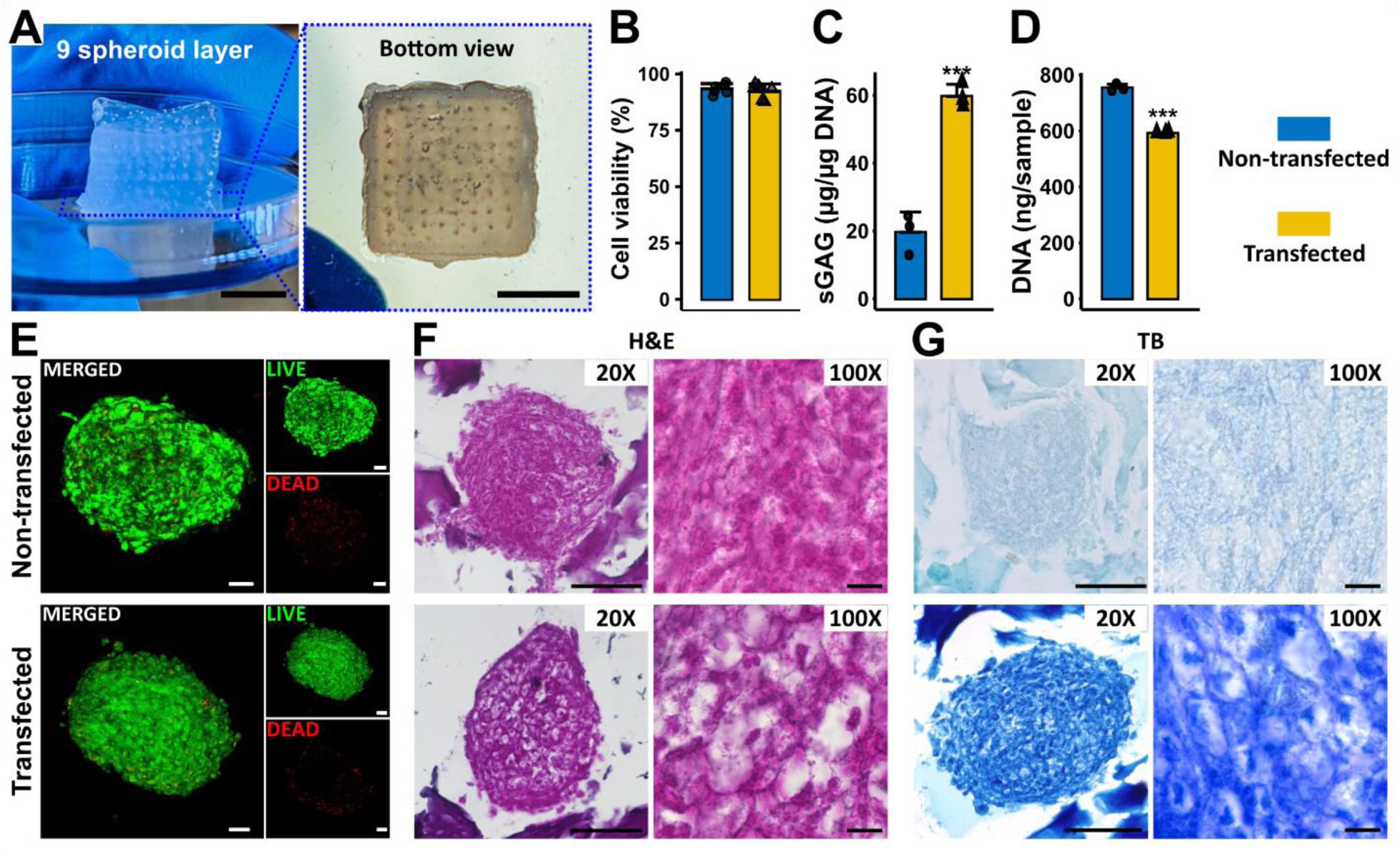
HITS-Bio for SCT fabrication. (A) 1 cm^3^ of cartilage tissue with CARink was bioprinted with 576 spheroids. Scale bar: 5 mm. Comparison between non-transfected and co-transfected spheroids cultured in 1 cm^3^ tissue at Week 2 for (B) cell viability measurements (*n* = 6), (C) sGAG normalized with DNA content (*n* = 3, \*\*\**p* < 0.001), and (D) the total DNA content of tissue constructs (*n* = 3, \*\*\**p* < 0.001). (E) LIVE/DEAD staining of non-transfected and transfected spheroids at Week 2 (Scale bar: 50 µm). Histomorphometric characterization of sectioned SCT stained with (F) H&E with 20X and 100X magnification and (G) TB-stained images captured with 20X and 100X magnification (Scale bar: 100 µm (20X) and 10 µm (100X)).

## 3. Discussion

Current bioprinting techniques face critical challenges, such as achieving physiologically-relevant cell densities, enhancing throughput for scalable tissue fabrication, developing bioinks tailored for specific applications, and enabling in situ fabrication capabilities. Towards this, the pursuit of high-throughput bioprinting marks a vital development in achieving rapid and efficient tissue fabrication, driven by automation. These advancements are essential for meeting the increasing demand for complex tissue constructs that replicate native tissue architecture and function. In this study, a novel High-throughput Integrated Tissue Fabrication System for Bioprinting, termed "HITS-Bio," comprising four key components was developed. These include (i) software for customized control, (ii) digitally-controlled nozzle array (DCNA) facilitating precise and multiple spheroid placement, (iii) compatible bioinks (BONink and CARink) ensuring smooth extrusion and printability, and (iv) miRNA transfected osteogenic and chondrogenic spheroids for *de novo* tissue fabrication. The platform was assessed for its applicability in in-situ osteogenesis of bone tissue and scalability by fabrication of volumetric cartilage tissues. The developed HITS-Bio platform was operated by a custom-made hardware-software interface as shown in **Figure S2** and **Supplementary Video 1**. The automation control was executed via the LabView Software Control Panel, managing motion stages with high precision (∼1 µm in X, Y, and Z axis). DCNA allowed customization, where selective nozzles could be activated depending on the target design. Initial testing of HITS-Bio was performed using 16 spheroids, which were selectively picked and placed alternately between red and blue dyed spheroids (Figure 2I). The process was completed under 30 sec, representing a significant speed compared to the existing benchmark in the literature ^9^, which required nearly 30 min.

Spheroids are promising candidates as building blocks for tissue fabrication as they recapitulate the native tissue environment with similar cell density and ECM composition and have the potential to rapidly induce tissue regeneration due to initially-delivered large pre-committed cell numbers ^9^. When spheroids are loaded in gels, they show better cell spreading and proliferation, and tissue-specific differentiation, compared to conventional cell-laden hydrogels ^18^. Until now, spheroid bioprinting techniques enabling the deposition of spheroids onto a gel substrate have been mainly limited to extrusion-based bioprinting (EBB) ^19^ and AAB ^9^. EBB with spheroids has been used to overcome the limitation of increasing cell density in an extrudable bioink ^5,20^. However, spheroids are randomly dispersed in the bioink, making their controlled positioning a challenge, resulting in area-to-area and batch-to-batch inconsistencies and limitations in loading efficiency in bioprinted constructs, similar to the case shown in **Figure S5**. To overcome these limitations, AAB has been utilized for high-precision bioprinting, where introduced, various tissue complexes have been fabricated inside functional hydrogels as well as support baths depending on the application ^12^. However, in these strategies, the placement of only one spheroid at a time was feasible, rendering scalable tissue fabrication a challenging task. Alternatively, multiple studies have demonstrated the potential of multi-nozzle bioprinting for rapid tissue fabrication ^21,22^. For example, a study by Hansen *et al.* used a multi-nozzle array to produce a hierarchically branched, microvascular network and exhibited high-throughput printing of single and multiple extrudable inks over large areas (1 m^2^). The strategy resulted in a significant reduction in printing time, where a 3D construct that takes a day to print using a single nozzle printhead took only 22 min to print using a system with 64 nozzles ^23^. However, to demonstrate large-scale patterning, the study used wax as an ink to print onto a 1-m^2^ glass substrate using the 64-nozzle printhead. In the current work, HITS-Bio was used to accelerate the spatial positioning of spheroids via their simultaneous deposition onto a gel substrate for scalable tissue fabrication. Spatial positioning of spheroids is important for mimicking native tissue microenvironments and promoting effective cell-cell communications, ensuring tissue functionality and organization ^4^. Herein, DCNA enabled the spatial arrangement of miR transfected spheroids. A combination of miR-196a-5p and 21 was used for the co-transfection of hADSCs to create osteogenically-committed spheroids. miR-196a-5p plays a crucial role in bone homeostasis and is highly expressed in osteoclast precursors ^24^. Kim *et al.* reported that miR-196a-5p regulates the proliferation and osteogenic differentiation of human ADSCs, which may be mediated through HOXC8 ^25^. Concurrently, miR-21 has been proven to play a role in bone formation by mediating mesenchymal stem cell proliferation and differentiation ^26,27^. It activates the ERK-MAPK (extracellular signal-regulated kinases (ERKs)-mitogen-activated protein kinases (MAPKs)) signaling pathway, promoting osteogenesis by suppressing the expression of its target gene SPRY1 ^28^. In a study, when combined, miR-196a-5p and -21 exhibit synergistic effects to enhanced osteogenesis, where miR-196a-5p stimulates osteogenic ability, while miR-21 further supports osteoblastic differentiation and amplified proliferation rate, confirming the hypothesis of Abu-laban *et al*. ^29^ and Celik *et al.* ^17^. Thus, spheroids with ∼8k cells/spheroid were formed in the current work and maintained a week in growth medium *in vitro*. After a week, spheroids reached

∼350 µm in diameter with sufficient structural properties for bioprinting purposes. These spheroids were then used along with the developed BONink for bone bioprinting.

This study introduced novel bioinks, characterized by their paste-like shear-thinning property enabling both *in vitro* and *in vivo* EBB. Bioink comprises GM (10 or 20% w/v), β-GP, HyA, fibrinogen, HA (15 or 30% w/v), and the photocrosslinker LAP. GM and HA served as bulk polymers influencing the bioink’s properties, while HyA and fibrinogen mimic ECM, promoting cell growth and tissue regeneration. β-GP enhances osteogenesis and mineralization, making these polymers ideal for bone regeneration applications ^30,31^. The developed bioinks were characterized rheologically, where flow sweep results indicated that the bioinks had shear-thinning attributes meeting the basic requirement for EBB. The decrease in viscosity upon increasing the shear rate implies a decrease in the extrusion pressure, facilitating smooth extrusion through smaller nozzles ^32^. Additionally, the bioinks possessed self-healing capability, which is essential for maintaining the integrity of bioprinted constructs. Self-healing shear-thinning bioinks stand out as promising materials for EBB ^33^. These bioinks can be extruded as their viscosity decreases under the shear, and subsequently self-heal once the shear is removed. This dual property ensures safe bioprinting of cells and the maintenance of shape fidelity post-bioprinting ^33^. Further, the assessment of mechanical properties of bioinks showed that the modulus falls within the range observed in native trabecular or cancellous bone, which typically exhibits elastic moduli of 0.02–2 GPa ^34^. Additionally, construct degradation at physiological conditions is considered advantageous because it allows for the construct to diminish so that the new ECM can slowly replace the degraded portions of the construct. The results indicated that an increase in the concentration of GelMA resulted in slower degradation, which may be attributable to increased methacrylamide crosslinks ^35^. The addition of HA and other components reduced the degradation rate further. This aligns with a prior study by Allen *et al.*, where a GelMA-gelatin-HA bioink was utilized for 3D bioprinting of bone constructs and their degradation with Type IV collagenase was significantly reduced by the addition of HA in a concentration-dependent manner ^36^. Based on these results, GM20HA30 composite was selected as a base cement material and termed ‘BONink,’ which was found to be biocompatible, affirming its suitability for bone tissue engineering applications.

The *in vitro* assessment results suggested that bioprinted constructs containing spheroids co-transfected with miR-(196a-5p + 21) showed superior upregulation of osteogenic genes. Interestingly, the expression of osteogenic markers may be influenced by the spheroid density. Early-stage markers like *RUNX2* and *BMP-4* show higher expression in low-density groups, likely due to better nutrient diffusion and efficient paracrine signalling, where factors can diffuse more evenly, promoting early differentiation. Conversely, high-density spheroids exhibit increased expression of intermediate to late-stage markers, such as *OSTERIX* and *BSP*, indicating enhanced maturation and mineralization, which may be due to closer cell-cell interactions and higher local concentrations of paracrine factors. These results suggest the role of spheroid density in optimizing osteogenic differentiation through paracrine signalling and mechanical cues. IHC staining confirmed the expression patterns, particularly highlighting intense staining for RUNX2 in the low-density group and OSTERIX in the high-density group, aligning with gene expression profiles. To mimic the native tissue physiology, engineered tissues require optimum cellular density and microstructural complexity ^37^. Studies have shown that spheroids play a crucial role in enhancing osteogenic differentiation. For instance, spheroids facilitate cell-to-cell interactions, leading to increased osteogenic potential ^3,38^. Moreover, the upregulation of osteogenic markers like RUNX2, BSP, and OSTERIX in miR-transfected spheroids has been linked to enhanced osteogenic differentiation ^17^.

Meanwhile, IOB is transforming surgical procedures by offering *in situ* fabrication of patient-specific tissue constructs directly at the surgical site ^39^. It enables precision customization, minimizing infection risks through the elimination of pre-fabricated implants, and improving healing due to freshly bioprinted tissues, while eliminating storage and transportation concerns. IOB has been previously utilized in repairing calvarial bone defects via laser-based bioprinting (LBB), where nano-HA was combined with collagen and MSCs to serve as the ink and deposited directly onto the calvarial defects in mice, resulting in a significant increase in bone formation observed after 2 months ^40^. However, LBB method is challenged by its slow deposition of biomaterials into a defect site. Further, the complexity of the laser set-up and its impracticality in surgical settings can restrict its potential for clinical translation. In another study, IOB was used for the reconstruction of craniomaxillofacial (CMF) tissues, including bone, skin, and composite (hard/soft) tissues. The use of a hybrid IOB approach (EBB and droplet-based bioprinting) reconstituted hard/soft composite tissues in a stratified arrangement resulted in ≈80% skin wound closure in 10 days and 50% bone coverage area at Week 6 ^41^. In the current work, for the first time, bioprinting was performed intraoperatively using spheroids under surgical settings to repair rat calvarial defects. The results showed that the newly formed bone invaded from one defect edge to another (Figure 4B) with a bone coverage area of 91 and 96% in 3 and 6 weeks for the high-density group, respectively. This demonstrates substantial bone regeneration in a short timeframe, attributed mainly to the innovative use of IOB with miR-transfected spheroids, which is quite challenging using other similar methods and materials. The maximum intensity projections from µCT data, used to score bony bridging, revealed significantly higher scores (3.85 ± 0.37 out of 4) for spheroid involved groups indicating near-complete bridging and enhanced bone regeneration, likely due to the superior osteogenic properties imparted by the miR transfection and BONink (Figure 4C). Moreover, the regenerated bone in this group exhibited significantly enhanced shear yield strength and modulus of resilience (Figure 4D). Histological analysis depicted the connectivity and compactness of the newly formed bone tissue with MT staining revealing dense blue islands indicative of immature bone tissue formation, particularly prominent in the high-density group. The study demonstrated superior regenerated bone *in situ*, supported by histological analysis showcasing dense, immature bone formation, which can be expected to develop into a mature bone in a longer timeframe *in vivo*. Overall, the findings support the feasibility of HITS-Bio as a powerful tool in IOB of spheroids for bone tissue with superior osteogenic potential, and high-throughput and speed, completing each construct in about 4.5 min. However, long-term assessment will be vital to establish the clinical relevance of these findings.

Furthermore, the HITS-Bio platform was explored for the scalability of bioprinted tissues. Thus, SCTs were successfully fabricated by depositing layers of CARink (representing GM20) using EBB followed by the precise placement of chondrogenic spheroids using DCNA, creating constructs with nine stacked tissue layers, comprising around 600 spheroids per SCT, demonstrating the potential for creating intricate, multi-layered tissues. The CARink, being transparent, allowed real-time monitoring of deposited spheroids with favorable extrusion properties (inset images in Figures 2Fii and **2Fiii**, and **Supplementary Video 7**). A combination of miR-140 and -21 was used for co-transfection of hADSCs to create chondrogenically-committed spheroids as per a previous report ^17^. Importantly, the results revealed that the process of transfection and tissue fabrication did not compromise spheroid viability (> 90%) compared to non-transfected spheroids. The dual miR-transfected SCTs exhibited a significantly higher sGAG content compared to the non-transfected control group (Figure 5C). sGAGs play a critical role in cartilage function, providing mechanical support and maintaining tissue hydration ^42^. The results suggest that HITS-Bio supported enhanced ECM deposition that is essential for cartilage formation. Interestingly, the DNA content was significantly lower in the transfected samples compared to the non-transfected samples (Figure 5D). This might be due to miR suppressing genes associated with stemness and promoting chondrocyte-specific markers and by downregulating cyclins and cyclin-dependent kinases, they limit cell proliferation, favoring differentiation ^25^. However, further investigation is warranted to understand the underlying mechanisms. Histological assessment using H&E confirmed the presence of chondrogenic spheroids within bioprinted constructs, where spheroids exhibited intense staining, indicating a higher density of ECM deposition with a developed lacunae-like structure (Figure 5F). Further, TB staining supported the presented quantitative sGAG data, where transfected spheroids displayed intense TB staining, emphasizing enhanced sGAG production (Figure 5G). Overall, the results demonstrated the efficacy of HITS-Bio and its potential application for the rapid bioprinting of cartilage tissues. Using HITS-Bio, the tissue dimensions were scaled up from a volume of ∼30 mm^3^ (disc with a diameter of 5 mm and a height of 1.5 mm) involving 64 spheroids for bone tissue bioprinted in 4.5 min per construct, to a volume of 1 cm^3^ (cube of 1 × 1 × 1 cm), incorporating 576 spheroids for cartilage tissue bioprinted in under 40 min (including EBB of the CARink). The ∼10-fold increase in the bioprinting speed was achieved through the integration of the aforementioned factors in HITS-Bio. However, because of the thick tissue, hypoxia in SCTs is possible and needs to be investigated in further studies. Nevertheless, in cartilage, it has been reported that the hypoxia condition protects against cartilage loss by regulating *Wnt* signalling ^43^.

Still, HITS-Bio has some aspects, where its performance could be enhanced. First, DCNA may have a clogging issue that can delay the bioprinting time. The clogging issue arises from cell debris accumulating in the spheroid chamber. Typically, when a nozzle becomes clogged, it is cleared by blowing air through the affected nozzle. Second, all nozzles in DCNA should be positioned on a uniform plane. This alignment is crucial for accurately patterning spheroids, ensuring they are placed at the same level, otherwise, loaded spheroid may penetrate deeper or remain elevated, which could lead to potential damage or imprecise bioprinting due to insufficient contact with the surface. During DCNA manufacturing, the precision along the Z-axis is thus important. As shown in Figure 2C, although the groups showed no significant differences, a decrease in the inter-nozzle distance led to an increase in variation in Z-axis positional error due to the interference caused by tightly packed nozzles during the micromanufacturing process. The developed DCNA can be conveniently used to bioprint on flat surfaces; however, bioprinting on uneven surfaces like a non-planar defect area during IOB can pose challenges. Thus, it is important to level the defect area on a rat to match the bottom of the DCNA plane before initiating the HITS-Bio process either by angle adjusting the roll pitch yaw of DCNA or by adjusting the rat’s head angle parallel to the DCNA surface. Third, although we limited DCNA to 4 x 4 nozzles in the current study, we can reconfigure it (e.g., 10 × 10 nozzle array) to further expedite the bioprinting process. This would eliminate the periodicity constraints imposed by the current DCNA design while simultaneously leading to improved tissue fabrication efficiency and better capabilities for various applications. Fourth, the experienced capillary reaction results in liquid elevation between the nozzles of DCNA. We silicon-coated the nozzles to overcome this and observed that the elevated liquid decreased as the inter-nozzle distance increased. Lastly, during spheroid aspiration, a minimum pressure of ∼3 mmHg was sustained throughout the whole DCNA, preventing media leakage when spheroids were placed with closed channels. It was noted that leaked media on the substrate hindered the placement of spheroids or caused them to move, resulting in inaccurate positioning. Importantly, achieving adequate hemostasis before the IOB process is essential, as bleeding during surgery may reduce accurate spheroid positioning. Thus, based on the potential to improve the aforementioned aspects of DCNA, the next generation of DCNA would ideally feature independently height-adjustable nozzles with an increased number of nozzles demonstrating multi-nozzle EBB for conformal bioprinting ^44^. Using the modified DCNA, HITS-Bio can be more accurate on adjustable surfaces and expand their prospective applications and freeform bioprinting of spheroids in support materials for complex shape fabrication.

The future research of this study aligns with the pressing need to advance spheroid bioprinting techniques toward the rapid fabrication of scalable vascularized tissues. Integrating vascular networks within large-scale bioprinted tissues, particularly for organs with high metabolic demands such as the heart, pancreas, or liver, signifies a crucial step towards achieving clinically-relevant volumes for transplantation. In addition, HITS-Bio can be utilized for developing *in vitro* models for drug screening or to understand interactions of multiple types of tissues using not only precisely bioprinted spheroids but also organoids at a controlled distance apart.

## 4. Conclusion

This study introduces HITS-Bio, a high-throughput bioprinting platform, that enables scalable tissue fabrication by precisely positioning a number of spheroids at an unprecedented speed using a digitally-controlled nozzle array (DCNA) platform. With a significant increase in the bioprinting performance compared to conventional techniques, HITS-Bio enabled the rapid creation of tissue via accelerated spatial positioning of spheroids on a gel substrate. The performance of HITS-Bio was exemplified by bioprinting multiple tissue types including bone and cartilage. The study showed for the first time intraoperative bioprinting with spheroids, where miR co-transfection enhanced osteogenesis in committed spheroids derived from hADSCs, demonstrating significant potential in repairing rat calvarial defects. Moreover, the potential of HITS-Bio for scalable tissue fabrication was demonstrated by the fabrication of scalable cartilage constructs with an unprecedented speed (under 40 min per construct) that surpasses the capabilities of conventional bioprinting technologies. HITS-Bio’s versatility showcases significant advancement in fabrication performance, and with the integration of IOB, it has the potential for personalized tissue biofabrication.

## 5. Methods

### 5.1 Cell culture and spheroid fabrication

Human adipose-derived stem cells (hADSCs, PT-5006, Lonza, Walkersville, MD) were obtained and cultured in a basal medium consisting of a 1:1 mixture of HyClone Dulbecco’s modified Eagle medium (F12) (Hyclone, Marlborough, MA) supplemented with 20% fetal bovine serum (FBS, R&D Systems, Minneapolis, MN), 1% penicillin/streptomycin (P/S, Corning, Manassas, VA) at 37 °C with 5% CO_2_ in a humidified incubator. The cell medium was changed every other day. hADSCs from passages 1–3 were used in experiments. For spheroid fabrication, the expanded hADSCs were trypsinized, centrifuged, and transferred to each well of a U-bottom 96-well plate (Greiner Bio One, Monroe, NC) with 7,000 cells per well to obtain spheroid clusters of 300-350 µm, followed by their incubation in a humidified incubator with 5% CO_2_ at 37 °C overnight allowing the spheroid formation as described in a previous work ^17^.

### 5.2 Development of the HITS-Bio platform

The HITS-Bio platform integrated X, Y, and Z motion axis for 3D movement, equipped with cameras, a digitally controllable nozzle array (DCNA) consisting of 30G nozzles and associated controls such as pressure regulators, flow controllers, and positioning systems (**Figure S2**). In the bioprinting stage, two 35-mm Petri dish were accommodated to be used as spheroid reservoir and tissue fabrication area, respectively. Solenoid valves and a pressure sensor digitally regulated the aspiration force and positive pressure, allowing for selective control and internal pressure monitoring of each nozzle. The hardware component of DCNA consisted of stainless-steel needles (30G, an inner diameter of 150 µm and an outer diameter of 305 µm) arranged in a 4 × 4 array with predefined spaces between nozzles, which was enabled by precisely stacked multiple acrylic plates micro-manufactured by laser cutting (**Figure S1**). These nozzles were carefully inserted through the holes on the plates, and after calibrating the surface and arranging the nozzles on the same plane, were adhered to the acrylic plates for stability. The assembled DCNA was coated with Sigmacote^®^ (SL2, Sigma-Aldrich, MO) to siliconize the surface to reduce the surface tension generated between the cell medium and DCNA. The coated DCNA was attached to the HITS-Bio platform via an adaptor, facilitating exact roll-pitch-yaw adjustments to align the DCNA parallel to the bioprinting workspace. An extrusion head for bioink deposition was integrated into DCNA (**Figure S2B**), with real-time bioprinting visualization and positional verification of DCNA and extrusion nozzle provided by three microscopic cameras offering isometric, bottom, and side views. Vacuum chambers prevented liquid from damaging the solenoid valves during the HITS-Bio process. The entire platform operated through a custom hardware-software interface (**Figure S2B** and **Supplementary Video 1**), enabling precise operation of motion stages, digital control of solenoid valves, monitoring and recording via a computer vision system, and real-time pressure value monitoring within the pneumatic line through a pressure sensor. Defined positions were saved as a .CSV file for reuse, and G-code-derived motion paths were uploaded for automated bioprinting. The system supported 2D and 3D visualization and analysis of objects for position, quantity, shape, and interspace. It also tracks their movement and incorporated safety features like an emergency stop button and maximum axis limits. The bioprinting process, standardized at 10 mm/s for spheroids 300-350 µm in diameter, was adjusted to 0.2 mm/s for lifting spheroids from the spheroid plates to air. Various DCNA sizes, designed for specific nozzle array widths and spacing, were fabricated, and assessed for positional accuracy and angular deviations, potentially resulting from handling errors during micro-manufacturing. Spheroid lifting percent success rates (*SR*%) for different DCNAs were calculated using the formula:

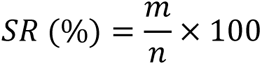

where *m* is the number of successfully lifted spheroids using DCNA and *n* is the total nozzle count with the platform designed for a 4 × 4 nozzle array, making *n* = 16. The HITS-Bio platform’s efficacy was benchmarked against a conventional single-nozzle AAB technique ^13,45,46^ using the same bioprinting parameters. Bioprinting time was documented for sets of 16, 32, and 64 spheroids. The platform’s versatility was further demonstrated using three different diameters of spheroids (∼350 μm stained with phalloidin AF488 (green), ∼425 μm stained with DAPI (blue) and ∼500 μm stained with phalloidin AF647 (red)) deposited on a 10% GelMA (GM10) substrate, as prepared below.

### 5.3 Preparation of Bioinks (BONink and CARink)

Gelatin methacryloyl (GM) was synthesized following a previously established protocol ^47,48^. GM was then reconstituted into 5 and 10% (w/v) solutions using warm PBS and incubated at 37 °C until fully dissolved. For preparation of the cartilage ink (CARink) (**Figure S6B**), the GM solution was combined with 5 mg/mL fibrinogen (F8630, Sigma-Aldrich, MO), 3 mg/mL hyaluronic acid sodium salt (HyA, 53747, Sigma-Aldrich, MO), and 0.25% (w/v) lithium phenyl (2,4,6-trimethylbenzoyl) phosphinate (LAP, L0290, TCI Chemicals, OR) as a photoinitiator. Concurrently, the bone ink (BONink) (**Figure S6A**) was formulated with the same components as CARink, along with the addition of 15 mg/mL β-glycerophosphate disodium salt hydrate (β-GP, G9422, Sigma-Aldrich, MO) and hydroxyapatite (HA, nanoXIM•HAp202, Fluidinova, Maia, Portugal) at 15 and 30% (w/v) concentrations, having a median particle size (d_50_) of 5.0 ± 1.0 μm. Both bioinks were homogenized using a FlackTek SpeedMixer (Louisville, CO) at 2,000 rpm for 3 min, followed by a subsequent 3-min cycle at the same speed to eliminate air bubbles. The resulting materials were carefully transferred into 1 cc syringes, avoiding the incorporation of air, and then wrapped in aluminum foil to shield from light-induced photo-crosslinking. They were stored vertically at room temperature (RT) for 2 h before bioprinting. After bioprinting with the extrusion of CARink and BONink, 25 U/mL of thrombin (T4648, Sigma-Aldrich, MO) was used with the cell culture medium as an enzymatic crosslinker for fibrinogen. The concentrations of the composite materials in BONink (GM and HA) and CARink (GM) were optimized for extrudability, mechanical stability, rheological properties, and biodegradability as described below.

### 5.4 Bioink characterization

#### 5.4.1 Rheological properties

Rheological properties were characterized using a rheometer (MCR 302, Anton Paar, Austria). All rheological measurements were performed in triplicates with a 25 mm parallel-plate geometry at RT (22 ℃) after having 2 h of resting at RT. The frequency sweep test was carried out in an angular frequency range of 0.1-100 rad/s at a strain in the linear viscoelastic region to examine the storage (G’) and loss (G") modulus without destroying the sample. To investigate the shear-thinning behavior of prepared bioinks, a flow sweep test was conducted in the shear rate range of 0.1-100 s^-1^. A recovery sweep test was also performed to evaluate changes in the viscosity after applying low and high shear rates at five intervals: (1) a shear rate of 0.1 s^-1^ for 30 sec, (2) a shear rate of 100 s^-1^ for 10 sec, (3) a shear rate of 0.1 s^-1^ for 30 sec, (4) a shear rate of 100 s^-1^ for 10 sec, and (5) a shear rate of 0.1 s^-1^ for 30 sec.

#### 5.4.2 Compression testing of in vitro samples

Unconfined compression tests were conducted using an Instron 5966 series advanced electromechanical testing system with a 10 kN load cell, following American Society for Testing and Materials (ASTM) standard D395–18 guidelines. Briefly, 13 × 6 mm (diameter × height) cylindrical specimens were compressed at a rate of 1.3 mm/min until reaching failure or 70% strain of their original height at RT. The obtained data was converted to stress-strain and the initial bulk modulus (kPa) was calculated from the initial gradient of the resulting curve within the range of 0–10% compressive strain.

#### 5.4.3 Biodegradability characteristics

To study the degradation characteristics of the prepared bioinks, we used the traditional gravimetric approach ^49^. Briefly, the initial weight of constructs was recorded, followed by their immersion in PBS with a pH of 7.4 and PBS containing 1 U/mL of collagenase Type II (Thermo Fisher Scientific, Waltham, MA) at a temperature of 37 °C until they reached a state of equilibrium. The collagenase solution was refreshed every 2-3 days to maintain constant enzyme activity. The weight of constructs was measured at different time points (Days 0, 3, 5, 7, 10, and 14). The degradation study was performed in triplicates per each time point and the percent degradation (*D*%) was determined using the following equation:

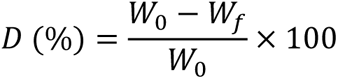

where, 𝑊_0_ and 𝑊_𝑓_ were the initial and measured weight of the construct at each time points, respectively.

#### 5.4.4 Biocompatibility

To evaluate the biocompatibility of BONink and CARink, hADSC spheroids were mixed into the bioinks and cultured at 37 °C in 5% CO_2_ in a humidified atmosphere. Cell viability in BONink was tested using an alamarBlue HS cell viability reagent (A50100, Thermo Fisher, MA) according to the manufacturer’s instructions on Days 1, 3, 7, and 14. Briefly, samples were treated with 10% (v/v) dye solution and incubated for 3 h. Next, 100 μL of the culture medium was analyzed using a microplate reader (Tecan Infinite 200 Pro, Switzerland) at 570/600 nm (excitation/emission). The results were expressed as the normalized value of the reduced dye, which correlated with the quantity of viable cells.

### 5.5 MicroRNA transfection for osteogenic and chondrogenic differentiation

MicroRNA (miR) mimics were used for osteogenic and chondrogenic differentiation according to the protocol reported in a previous work ^17,50^. Custom oligonucleotides (miR-140: 5’-CAGUGGUUUUACCCUAUGGUAG-3’; miR-21: 5’CAACAGCAGUCGAUGGGCUGU 3’; miR-196a-5p: 5’-UAGGUAGUUUCAUGUUGUUGGG-3’) were ordered from Integrated DNA Technologies (Coralville, IA). As described in a previous work ^17^, transfection was performed using lipofectamine RNAiMAX transfection reagent (Thermo Fisher Scientific, Waltham, MA), where lipofectamine was mixed with miR mimics according to the manufacturer’s protocol. hADSCs were co-transfected with miR-(196a-5p + 21) mimic for osteogenic differentiation and miR-(140 + 21) mimic for chondrogenic differentiation before being seeded in Opti-MEM medium (Gibco, Carlsbad, CA) in 175 cm^2^ cell culture flasks for 24 h. The final concentration of all miR transfections in the opti-MEM medium was determined to be 200 nM for a volume of 10 mL. The 10 mL solution with a final concentration of 200 nM was transferred to 175 cm^2^ cell culture flasks and incubated at 37 °C and 5% CO_2_ for 24 h. Transfected cells were collected by trypsinization and formed as spheroids using the protocol explained in Section 5.1. After transfection, a basal medium, consisting of a 1:1 mixture of HyClone Dulbecco’s modified Eagle medium (F12) supplemented with 2% FBS, and 1% P/S, was used for miR-transfected spheroids. A commercially available osteogenic differentiation medium (Catalog number: 417D-250, Cell Applications, San Diego, CA) was used as a positive control group for osteogenic differentiation.

### 5.6 In vitro bioprinting process for bone tissue fabrication

Bone constructs were fabricated using osteogenically-committed spheroids and the BONink in a stratified manner utilizing a hybrid approach. First, the BONink was extruded into a disk shape of 5 mm diameter with a 22G tapered tip (inner diameter of 410 µm) using 80 kPa pneumatic pressure. Subsequently, osteogenic spheroids were precisely bioprinted atop the BONink layer, with densities categorized as 16 spheroids for low density and 64 spheroids for high density. Another layer of BONink was then extruded over the bioprinted spheroids to achieve a total thickness of ∼1.5 mm. This assembly was photo-crosslinked using a 405 nm near UV light source for 1 min. The bioprinting process was conducted at RT within a laminar flow biosafety cabinet to maintain sterility. Once bioprinted, the structures were cultured in 12-well plates with the basal or osteogenic differentiation media, incubated at 37 °C in a humidified 5% CO_2_ atmosphere, and the media refreshed every other day. Samples were collected at Weeks 2 and 4 for further characterization.

### 5.7 Intraoperative bioprinting of bone

For IOB aimed at bone regeneration in CMF defects, inbred immunodeficient RNU athymic rats (both male and female), acquired at 5 weeks of age from Charles River Laboratories International, Inc., Wilmington, MA, were cared for and aged at our animal facility until the age of 12 weeks. This care followed guidelines established by the American Association for Laboratory Animal Science (AALAS) and procedures approved by the Institutional Animal Care and Use Committee (IACUC, protocol #46591). A total of 14 rats were divided into four groups: empty defect (control), BONink only, low density, and high density. At Week 12, the rats underwent survival surgeries. Anesthesia was induced using Isoflurane in a concentration range between 2 and 5% delivered in oxygen at a flow rate of 0.5 to 2 L min^-1^. In addition, bupivacaine (0.015 mg kg^−1^) (Centralized Biological Laboratory, PSU) at a concentration of 2.5 mg mL^−1^ and buprenorphine (0.015 mg kg^−1^) were injected subcutaneously before the surgery. Once fully anesthetized, artificial tears were applied to the rats’ eyes and their heads were shaved and cleansed with betadine surgical scrub followed by ethanol. A sagittal incision, ∼2 cm long, was made on the skin and the periosteum was retracted to expose the calvarium. Two critical size calvaria defects, each 5 mm in diameter, were then created on either side of the parietal bone on the rat skull using a trephine bit, ensuring the Dura mater remained intact. The bone construct was bioprinted using HITS-Bio in a surgical setting (**Figure S2A**). The BONink was first extruded directly into the defect to create the base layer. Using HITS-Bio, osteogenically-committed spheroids were then bioprinted on this layer in either low (16 spheroids) or high (64 spheroids) densities. A final layer of the BONink was applied over the spheroids and crosslinked with 405 nm near UV light for 1 min. Post bioprinting, the skin was sutured with 4-0 vicryl sutures (Ethicon Inc., Somerville, NJ). Animals were placed on a warming pad for recovery. They were closely monitored until they regained sternal recumbency. Rats were observed daily and weighed for at least 10 days post-surgery, and weekly thereafter. At 3 weeks, rats were anesthetized and scanned using μCT in the live condition. At 6 weeks post-surgery, rats were euthanized by carbon dioxide (CO_2_) inhalation at a flow rate of 2 L min^-1^ until cessation of breathing, followed by decapitation for tissue collection. The collected tissues underwent histology and immunohistochemistry to evaluate calvarial bone regeneration.

### 5.8 Microcomputed tomography

μCT scanning was conducted to assess new bone formation, utilizing a vivaCT 40 scanner (Scanco Medical, Brüttisellen, Switzerland) with parameters set at 17.5 µm isometric voxels, 70 kV energy, 114 µA intensity, and 200 ms integration time, as detailed in a previously published work ^41^. Rats at Week 3 were anesthetized and scanned in the live condition as described above, while at Week 6 they were euthanized and scanned. Post-scanning, the data were refined and analyzed using Avizo software (FEI Company, Hillsboro, OR) for quantifying bone volume to total volume ratio (BV/TV (%)), normalized bone mineral density (BMD %), and bone coverage area (%). Importantly, the volume fraction of hydroxyapatite in the BONink was pre-calculated before conducting *in vivo* studies to establish a baseline for mineralization and to mitigate potential artifacts in scanned samples, which was found to be minimal. Further, μCT data was used to assess the extent of bony bridging within the defect, which was scored according to the grading scale as described in a previous report ^51^. These scores were determined from maximum intensity projections generated from the μCT datasets in the CT-analyser software (Bruker, Billerica, MA).

### 5.9 Mechanical testing of calvarial explants

Unconfined compression tests were conducted using an Instron 5966 mechanical testing system (Instron, Norwood, MA,) with a 10 kN load cell, using a modified fixture as per a previous report ^52^. For the setup, the explants were aligned on a plate with a central hole matching the defects, ensuring an unobstructed path for the probe. Push-out testing was performed to assess the mechanical properties of bone specimens harvested at Week 6. Using a 4.5-mm diameter probe, uniaxial compression was applied to the bone specimens at a crosshead speed of 0.01 mm/s, pushing the probe through the defects and the hole in the plate to measure the force required for bone-implant dislodgement. The test was stopped when the probe passed through the defects. The obtained data was assessed, and the shear yield strength (MPa), modulus of resilience (MPa) and shear modulus (MPa) were calculated from the initial gradient of the resulting curve within the range of 0–10% compressive strain.

### 5.10 Histological assessment

*In vitro* bone tissue samples were fixed in 4% paraformaldehyde (PFA) overnight at RT at Weeks 2 and 4 post bioprinting. The fixed samples were dehydrated in a series of graded alcohol solutions and embedded in paraffin blocks using a Leica TP 1020 automatic tissue processor (Leica). Subsequently, 10 µm sections were cut using a Shandon Finesse Paraffin microtome (Thermo Electron Corporation, Waltham, MA) and transferred onto positively charged slides. *In vivo* calvarial tissue samples were harvested at Week 6 following µCT scanning. These tissues were washed with PBS and fixed in 10% PFA for 2 days. Post-fixation, samples were washed again with PBS and subjected to decalcification in 0.5 M ethylenediaminetetraacetic acid (EDTA) disodium salt solution (Research Products International, Mt. Prospect, IL) for 6 weeks. After decalcification, samples were embedded in O.C.T. cryomatrix (Thermo Fisher Scientific, Waltham, MA) and sectioned at 15 µm thickness using a Leica CM1950 cryostat (Leica, Buffalo Grove, IL) at -20 °C. The resultant sections were used for subsequent histology and immunofluorescence staining. For hematoxylin and eosin (H&E) staining and deparaffinization, a Leica Autostainer ST5010 XL was used. The H&E-stained sections were then mounted and visualized using a Zeiss Axiozoom V16 inverted fluorescence microscope (Zeiss, Oberkochen, Germany). To evaluate collagen deposition, Masson’s Trichrome (MT) staining was performed following the manufacturer’s protocol (Sigma-Aldrich). After dehydration, the sections were mounted and imaged with a Keyence BZ-9000 microscope (Keyence Corporation of America, Elmwood Park, NJ).

### 5.11 Immunohistochemistry

For immunohistochemical (IHC) staining, sections from paraffin-embedded samples underwent deparaffinization while cryo-sectioned samples were thawed for 10–20 min at RT from -30 °C. All sections were permeabilized using 0.2% Triton X-100 for 10 min, washed in 1X PBS, and blocked with 2.5% normal goat serum (NGS) for 60 min at RT to prevent non-specific binding. To detect the bone tissue, sections were incubated with a mouse anti-RUNX2 (runt-related transcription factor 2) primary antibody (1:20 in 2.5% NGS, Cat. No. ab76956; Abcam) and a rabbit anti-Sp7/Osterix (OSTERIX) primary antibody (1:100 in 2.5% NGS; Cat. No. ab209484, Abcam) or a rabbit anti-bone sialoprotein (BSP) primary antibody (1:100 in 2.5% NGS; Cat. No. ab52128, Abcam) overnight. After washing with PBS, sections were incubated with goat anti-mouse IgG (H + L) Alexa Fluor 488 and goat anti-rabbit IgG (H + L) Alexa Fluor 647 secondary antibodies (1:200 in PBS; Cat. Nos. A11017 and A21245, Invitrogen) for 3 h. Following additional washes, the sections were mounted with ProlongTM Gold antifade reagent containing DAPI and imaged using an LSM 880 Zeiss confocal microscope (Zeiss, Oberkochen, Germany). To visualize the cartilage tissue, sections were similarly processed and incubated with mouse anti-Aggrecan (BC-3) primary antibody (1:50 in 2.5% NGS; Cat. No. MA3-16888; Thermo Fisher Scientific) and rabbit anti-Col-II primary antibody (1:100 in 2.5% NGS; Cat. No. ab34712, Abcam) overnight, followed by respective secondary antibodies and imaged using the Zeiss LSM 880 confocal microscope.

### 5.12 Gene expression using quantitative real-time polymerase chain reaction (qRT-PCR)

To assess osteogenic gene expression profiles through quantitative real-time PCR (qRT-PCR), RNA was isolated from each sample using TRIzol reagent (Life Technologies, Carlsbad, CA) and subsequently purified using the RNA Mini Kit (Thermo Fisher Scientific, Waltham, MA) as per the manufacturer’s protocol at Weeks 2 and 4. The concentration of RNA from each sample was determined by measuring the absorbance at a ratio of 260/280 nm using a Nanodrop (ND-1000, Thermo Fisher Scientific, Waltham, MA). The isolated RNA was then converted to cDNA using AccuPower^®^ CycleScript RT PreMix (BIONEER, Korea) following the manufacturer’s guidelines. Quantitative analysis of gene expression was performed using PowerUp™ SYBR™ Green Master Mix (Thermo Fisher Scientific, Waltham, MA) on a Quant Studio 3 PCR system (Thermo Fisher Scientific, Waltham, MA). The set of osteogenic genes analyzed included collagen type-1 (*COL-1*), runt-related transcription factor-2 (*RUNX2*), bone sialoprotein (*BSP*), bone morphogenetic protein-4 (*BMP-4*), and transcription factor Sp7 (*OSTERIX*). The primer details for these genes were provided in **Table S2**. The fold-change in gene expression of the target genes was quantified using the 2^−ΔΔCT^ method and then normalized to glyceraldehyde 3-phosphate dehydrogenase (*GAPDH*) as a housekeeping gene. The fold change of negative control (non-transfected hADSCs) on Day 1 was set as 1-fold and values in all groups were normalized to that of the group.

### 5.13 Scalable cartilage tissue fabrication

For the scalable cartilage tissue (SCT) fabrication, CARink and chondrogenic-committed transfected spheroids were alternately bioprinted to create a 1 cm³ construct. A week after the transfection process, spheroids were deposited onto 1 × 1 cm square extruded CARink. The deposition of spheroids and CARink was repeated nine times, layering 64 spheroids into each layer to accumulate 576 spheroids per sample in a stratified manner, resulting in a 1 cm³ cartilage tissue construct. The constructs were then photocrosslinked for 3 min and cultured in basal medium for two weeks *in vitro* to assess cartilage development.

### 5.14 Cell viability of CARink

LIVE/DEAD staining was performed to assess the cell viability of CARink. The fabricated scalable cartilage tissue was collected on Day 14 and washed with PBS, stained with a working solution composed of 2 µM calcein AM (Invitrogen, CA) and 4 µM ethidium homodimer-1 (EthD-1, Invitrogen, CA) for 45 min in an incubator and then imaged using the LSM 880 Zeiss confocal microscope. Cell viability (%) was determined after the deconvolution process to reduce the background signal based on the number of cells that had green or red fluorescence signal by dividing the number of green-fluorescent cells by the total number of cells and multiplying by 100. Image J software (National Institutes of Health, MD) was used for image analysis.

### 5.15 Physicochemical characterization of SCT

To quantity the sulfated glycosaminoglycan (sGAG) content of SCT, we examined both miR-(140 + 21) co-transfected and non-transfected hADSCs spheroids using a glycosaminoglycans assay kit (Chondrex, Inc., Redmond, WA). For each group, three samples were collected at Week 2 and then washed with PBS. Proteinase K (Sigma Aldrich, MO) was added to each sample at a final concentration of 0.5 mg/mL and incubated overnight at 56 °C for enzymatic cell lysis and DNA release. After incubation, 1,9-dimethylmethylene blue (DMMB) solution was added following the manufacturer’s protocol and the absorbance was measured at 525 nm using a spectrophotometer (Infinite 200 Pro, Tecan, Männedorf, Switzerland). The sGAG concentration was then normalized to the dsDNA content of each sample. DNA quantitation was performed using a Quant-iT PicoGreen dsDNA Assay Kit (Molecular Probes Inc., Eugene, OR) according to the manufacturer’s guidelines. Samples were excited at 480 nm and emission was measured at 520 nm using the spectrophotometer. The total DNA concentration was determined by comparison with a standard curve of Lambda DNA standard solution to ascertain DNA amounts. For histomorphometric analysis, SCT samples were fixed at Week 2 post bioprinting, dehydrated, and then sectioned. For H&E staining, resultant sections were stained using the Leica Autostainer ST5010 XL. The H&E-stained sections were then mounted and visualized using the Zeiss Axiozoom V16 inverted fluorescence microscope. For Toluidine blue O (TB) staining, sections were incubated in a 0.1% TB solution in deionized water at RT for 2 min. Following two rinses with deionized water, the sections were dehydrated using 95 and 100% ethanol, cleared with xylene, and mounted using Xylene Substitute Mountant (Epredia™, Kalamazoo, MI) before imaging with the Keyence BZ-9000 microscope.

### 5.16 Statistical analysis

All data are presented as mean ± standard deviation and analyzed by Minitab 17.3 (Minitab Inc., USA). Multiple comparisons were analyzed by one-way analysis of variance (ANOVA) followed by post-hoc Tukey’s multiple-comparison test to determine the individual differences among the groups. For comparisons between two different experimental groups, statistical significance was analyzed using two-sided t-tests. Statistical differences were considered significant at *p < 0.05, **p <0.01, ***p < 0.001.

## Acknowledgement

This work has been supported by National Institute of Biomedical Imaging and Bioengineering (NIBIB) Award R01EB034566 (I.T.O.) and National Institute of Dental and Craniofacial Research (NIDCR) Award R01DE028614 (I.T.O.). We thank Dr. Jian Yang (Penn State University) for providing access to material characterization equipment, Ethan M. Gerhard (Penn State) for his assistance in mechanical testing, Vaibhav Pal (Penn State) for his assistance with preparation of DCNA and GelMA, and Elisabeth G. Aliftiras (Penn State) for her assistance with illustrations.

## References

1 Murphy, S. V. & Atala, A. 3D bioprinting of tissues and organs. Nature biotechnology 32, 773–785 (2014).

2 Ozbolat, I. T. & Yu, Y. Bioprinting toward organ fabrication: challenges and future trends. IEEE Transactions on Biomedical Engineering 60, 691–699 (2013).

3 Banerjee, D. et al. Strategies for 3D bioprinting of spheroids: A comprehensive review. Biomaterials 291, 121881 (2022).

4 Kim, S.-j., et al. Spatially arranged encapsulation of stem cell spheroids within hydrogels for the regulation of spheroid fusion and cell migration. Acta Biomaterialia 142, 60–72 (2022).

5 Liu, Y. et al. hESCs-derived early vascular cell spheroids for cardiac tissue vascular engineering and myocardial infarction treatment. Advanced Science 9, 2104299 (2022).

6 Gutzweiler, L. et al. Large scale production and controlled deposition of single HUVEC spheroids for bioprinting applications. Biofabrication 9, 025027 (2017).

7 Moldovan, N. I., Hibino, N. & Nakayama, K. Principles of the Kenzan method for robotic cell spheroid-based three-dimensional bioprinting. Tissue Engineering Part B: Reviews 23, 237–244 (2017).

8 Blakely, A. M., Manning, K. L., Tripathi, A. & Morgan, J. R. Bio-pick, place, and perfuse: a new instrument for three-dimensional tissue engineering. Tissue Engineering Part C: Methods 21, 737–746 (2015).

9 Ayan, B. et al. Aspiration-assisted bioprinting for precise positioning of biologics. Science advances 6, eaaw5111 (2020).

10 Roth, J. G. et al. Spatially controlled construction of assembloids using bioprinting. Nature Communications 14, 4346 (2023).

11 Han, G. et al. 3D Printed Organisms Enabled by Aspiration-Assisted Adaptive Strategies. Advanced Science **n/a**, 2404617 10.1002/advs.202404617

12 Ayan, B. et al. Aspiration-assisted freeform bioprinting of pre-fabricated tissue spheroids in a yield-stress gel. Communications physics 3, 183 (2020).

13 Heo, D. N. et al. Aspiration-assisted bioprinting of co-cultured osteogenic spheroids for bone tissue engineering. Biofabrication 13, 015013 (2020).

14 Daly, A. C., Davidson, M. D. & Burdick, J. A. 3D bioprinting of high cell-density heterogeneous tissue models through spheroid fusion within self-healing hydrogels. Nature communications 12, 753 (2021).

15 Utama, R. H. et al. A 3D bioprinter specifically designed for the high-throughput production of matrix-embedded multicellular spheroids. Iscience 23 (2020).

16 Salhotra, A., Shah, H. N., Levi, B. & Longaker, M. T. Mechanisms of bone development and repair. Nature reviews Molecular cell biology 21, 696–711 (2020).

17 Celik, N., Kim, M. H., Yeo, M., Kamal, F., Hayes, D. J. & Ozbolat, I. T. miRNA induced 3D bioprinted-heterotypic osteochondral interface. Biofabrication 14, 044104 (2022).

18 Heo, D. N., Hospodiuk, M. & Ozbolat, I. T. Synergistic interplay between human MSCs and HUVECs in 3D spheroids laden in collagen/fibrin hydrogels for bone tissue engineering. Acta biomaterialia 95, 348–356 (2019).

19 Jakab, K., Neagu, A., Mironov, V., Markwald, R. R. & Forgacs, G. Engineering biological structures of prescribed shape using self-assembling multicellular systems. Proceedings of the National Academy of Sciences 101, 2864–2869 (2004).

20 Decarli, M. C. et al. Bioprinting of Stem Cell Spheroids Followed by Post-Printing Chondrogenic Differentiation for Cartilage Tissue Engineering. Advanced Healthcare Materials 12, 2203021 (2023).

21 Liu, W. et al. Rapid continuous multimaterial extrusion bioprinting. Advanced materials 29, 1604630 (2017).

22 Skylar-Scott, M. A., Mueller, J., Visser, C. W. & Lewis, J. A. Voxelated soft matter via multimaterial multinozzle 3D printing. Nature 575, 330–335 (2019).

23 Hansen, C. J. et al. High-throughput printing via microvascular multinozzle arrays. Advanced Materials 25, 96–102 (2013).

24 Takafuji, Y., Tatsumi, K., Kawao, N., Okada, K., Muratani, M. & Kaji, H. MicroRNA-196a-5p in extracellular vesicles secreted from myoblasts suppresses osteoclast-like cell formation in mouse cells. Calcified Tissue International 108, 364–376 (2021).

25 Kim, Y. J., Bae, S. W., Yu, S. S., Bae, Y. C. & Jung, J. S. miR-196a regulates proliferation and osteogenic differentiation in mesenchymal stem cells derived from human adipose tissue. Journal of Bone and Mineral Research 24, 816–825 (2009).

26 Chen, C., Liu, Y.-M., Fu, B.-L., Xu, L.-L. & Wang, B. MicroRNA-21: an emerging player in bone diseases. Frontiers in Pharmacology 12, 722804 (2021).

27 Yang, N. et al. Tumor necrosis factor α suppresses the mesenchymal stem cell osteogenesis promoter miR-21 in estrogen deficiency–induced osteoporosis. Journal of Bone and Mineral Research 28, 559–573 (2013).

28 Mei, Y. et al. miR-21 modulates the ERK–MAPK signaling pathway by regulating SPRY2 expression during human mesenchymal stem cell differentiation. Journal of cellular biochemistry 114, 1374–1384 (2013).

29 Abu-Laban, M., et al. Combinatorial delivery of miRNA-nanoparticle conjugates in human adipose stem cells for amplified osteogenesis. Small 15, 1902864 (2019).

30 Burdick, J. A. & Prestwich, G. D. Hyaluronic acid hydrogels for biomedical applications. Advanced materials 23, H41–H56 (2011).

31 Chung, C.-H., Golub, E. E., Forbes, E., Tokuoka, T. & Shapiro, I. M. Mechanism of action of β-glycerophosphate on bone cell mineralization. Calcified Tissue International 51, 305–311 (1992).

32 Schwab, A., Levato, R., D’Este, M., Piluso, S., Eglin, D. & Malda, J. Printability and shape fidelity of bioinks in 3D bioprinting. Chemical reviews 120, 11028–11055 (2020).

33 Karvinen, J. & Kellomäki, M. Design aspects and characterization of hydrogel-based bioinks for extrusion-based bioprinting. Bioprinting, e00274 (2023).

34 Wang, X. et al. Topological design and additive manufacturing of porous metals for bone scaffolds and orthopaedic implants: A review. Biomaterials 83, 127–141 (2016).

35 Zhao, X. et al. Photocrosslinkable gelatin hydrogel for epidermal tissue engineering. Advanced healthcare materials 5, 108–118 (2016).

36 Allen, N. B., Abar, B., Johnson, L., Burbano, J., Danilkowicz, R. M. & Adams, S. B. 3D-bioprinted GelMA-gelatin-hydroxyapatite osteoblast-laden composite hydrogels for bone tissue engineering. Bioprinting 26, e00196 (2022).

37 Skylar-Scott, M. A. et al. Biomanufacturing of organ-specific tissues with high cellular density and embedded vascular channels. Science advances 5, eaaw2459 (2019).

38 Griffin, K. H., Fok, S. W. & Kent Leach, J. Strategies to capitalize on cell spheroid therapeutic potential for tissue repair and disease modeling. npj Regenerative Medicine 7, 70 (2022).

39 Wu, Y., Ravnic, D. J. & Ozbolat, I. T. Intraoperative bioprinting: repairing tissues and organs in a surgical setting. Trends in biotechnology 38, 594–605 (2020).

40 Keriquel, V. et al. In situ printing of mesenchymal stromal cells, by laser-assisted bioprinting, for in vivo bone regeneration applications. Scientific reports 7, 1778 (2017).

41 Moncal, K. K. et al. Intra-operative bioprinting of hard, soft, and hard/soft composite tissues for craniomaxillofacial reconstruction. Advanced functional materials 31, 2010858 (2021).

42 Alcaide-Ruggiero, L., Cugat, R. & Domínguez, J. M. Proteoglycans in articular cartilage and their contribution to chondral injury and repair mechanisms. International Journal of Molecular Sciences 24, 10824 (2023).

43 Bouaziz, W. et al. Interaction of HIF1α and β-catenin inhibits matrix metalloproteinase 13 expression and prevents cartilage damage in mice. Proceedings of the National Academy of Sciences 113, 5453–5458 (2016).

44 Uzel, S. G., Weeks, R. D., Eriksson, M., Kokkinis, D. & Lewis, J. A. Multimaterial multinozzle adaptive 3D printing of soft materials. Advanced Materials Technologies 7, 2101710 (2022).

45 Kim, M. H., Banerjee, D., Celik, N. & Ozbolat, I. T. Aspiration-assisted freeform bioprinting of mesenchymal stem cell spheroids within alginate microgels. Biofabrication 14, 024103 (2022).

46 Ayan, B., Wu, Y., Karuppagounder, V., Kamal, F. & Ozbolat, I. T. Aspiration-assisted bioprinting of the osteochondral interface. Scientific reports 10, 13148 (2020).

47 Zhu, M., Wang, Y., Ferracci, G., Zheng, J., Cho, N.-J. & Lee, B. H. Gelatin methacryloyl and its hydrogels with an exceptional degree of controllability and batch-to-batch consistency. Scientific reports 9, 6863 (2019).

48 Pal, V. et al. High-throughput microgel biofabrication via air-assisted co-axial jetting for cell encapsulation, 3D bioprinting, and scaffolding applications. Biofabrication 15, 035001 (2023).

49 Kang, Y. et al. Intraoperative bioprinting of human adipose-derived stem cells and extra-cellular matrix induces hair follicle-like downgrowths and adipose tissue formation during full-thickness craniomaxillofacial skin reconstruction. Bioactive Materials 33, 114–128 (2024).

50 Celik, N., Kim, M. H., Hayes, D. J. & Ozbolat, I. T. miRNA induced co-differentiation and cross-talk of adipose tissue-derived progenitor cells for 3D heterotypic pre-vascularized bone formation. Biofabrication 13, 044107 (2021).

51 Patel, Z. S., Young, S., Tabata, Y., Jansen, J. A., Wong, M. E. & Mikos, A. G. Dual delivery of an angiogenic and an osteogenic growth factor for bone regeneration in a critical size defect model. Bone 43, 931–940 (2008).

52 Lawson, Z. T., Han, J., Saunders, W. B., Grunlan, M. A., Moreno, M. R. & Robbins, A. B. Methodology for performing biomechanical push-out tests for evaluating the osseointegration of calvarial defect repair in small animal models. MethodsX 8, 101541 (2021).

