## Supplementary Information for "High-Throughput Bioprinting of Spheroids for Scalable Tissue Fabrication"

### Supplementary Figures

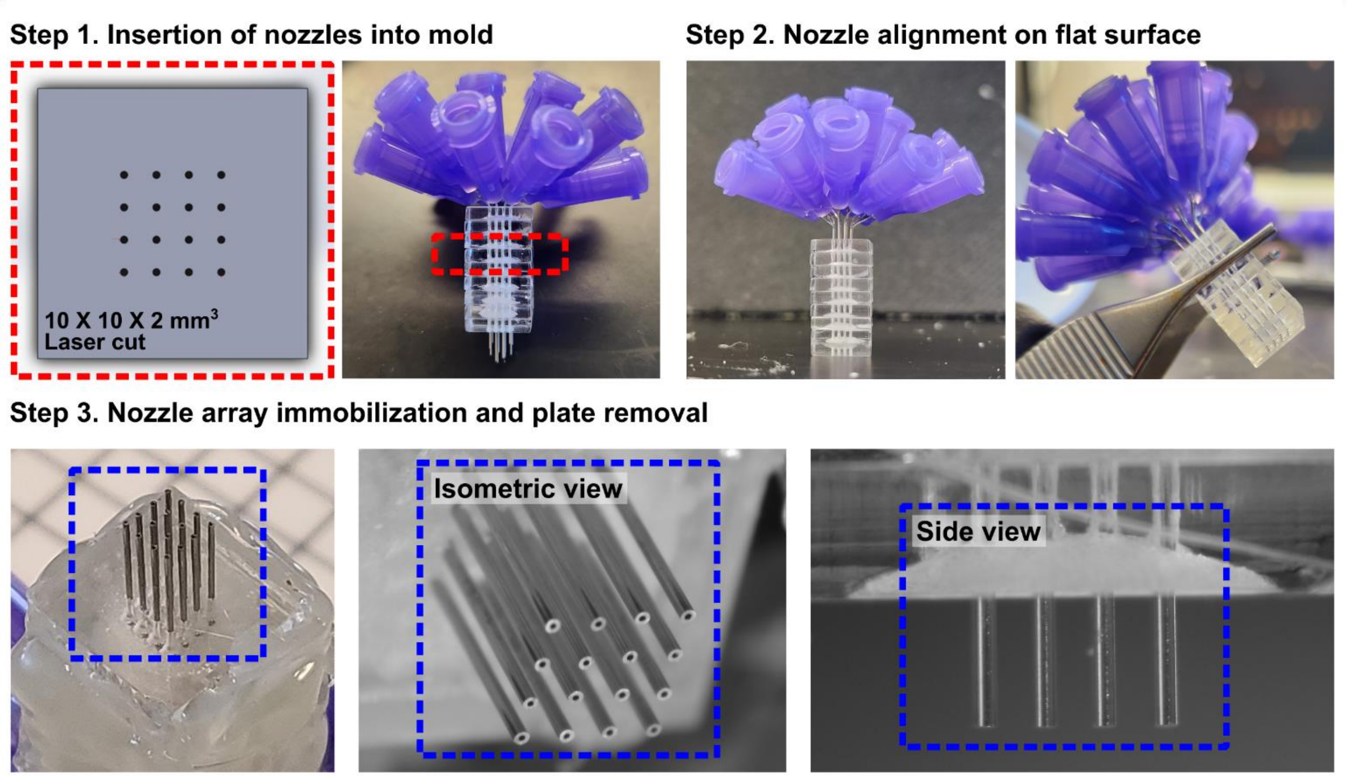


**Figure S1. Schematics illustrating the preparation of the DCNA nozzle.** First, nozzles were inserted into precisely stacked acrylic plates, micro-manufactured via laser cutting. Next, the nozzles were levelled and aligned by pressing them against a flat surface. Adhesive was then applied to immobilize the assembled nozzles. Finally, the two bottom acrylic plates were removed to obtain DCNA.

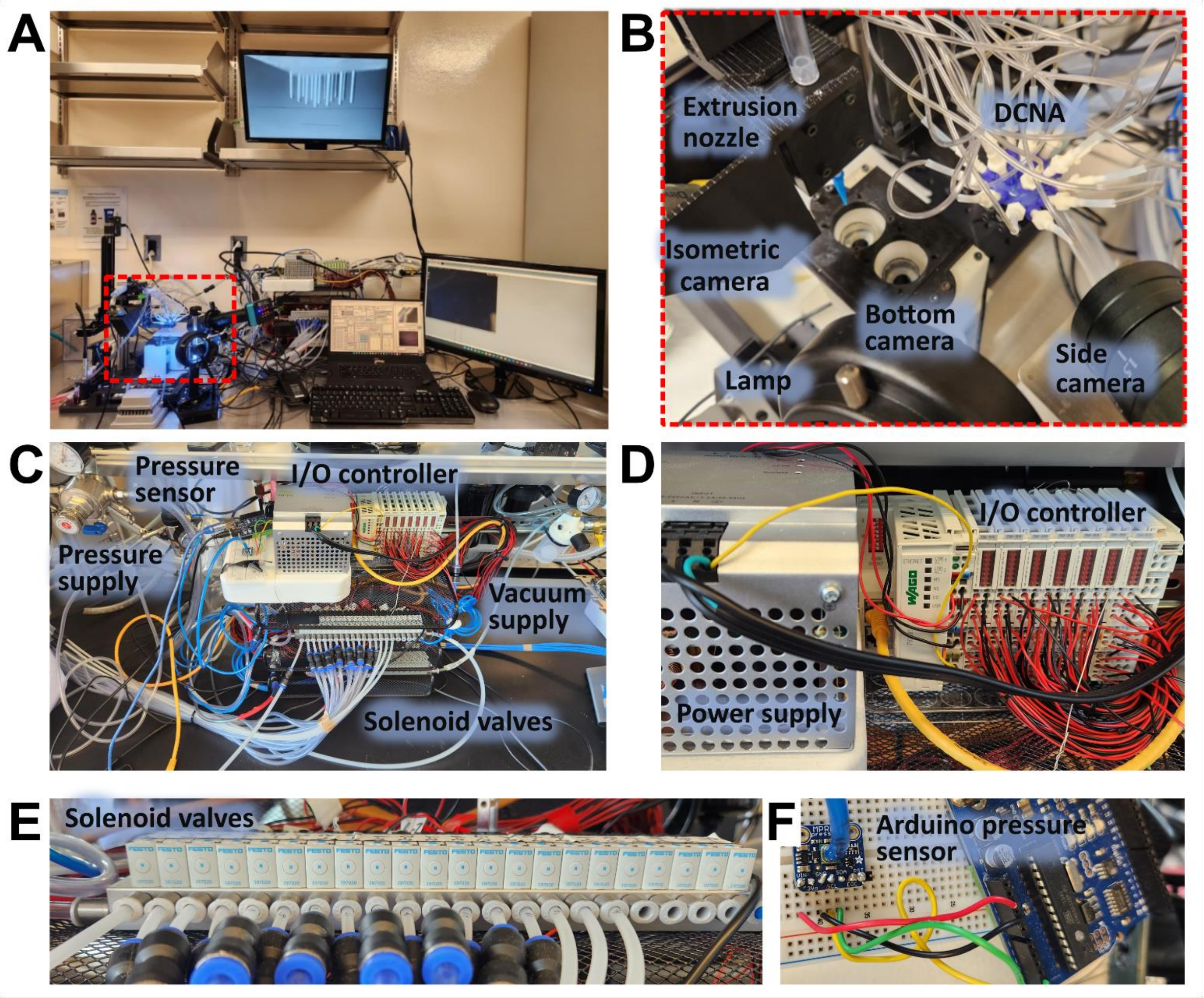


**Figure S2.** HITS-Bio hardware setup for (A) intraoperative bioprinting in a surgical setting. (B) HITS-Bio was composed of DCNA associated with an extrusion nozzle, cameras for isometric, side, and bottom views. (C) The components involved in pressure control, including the pressure sensor, pressure supply, vacuum supply, and solenoid valves, used for regulation of pressure/vacuum. (D) Input/output (I/O) controller and power supply, which manage the electronic control aspects of the system. (E) Solenoid valves, essential for regulating the pressure and vacuum in DCNA. (F) Arduino pressure sensor setup, utilized for real-time monitoring of pressure within the system.

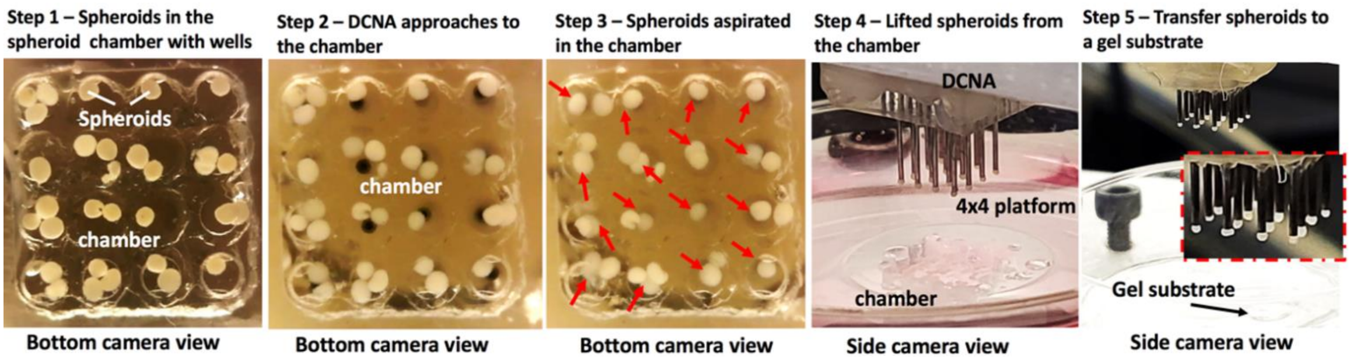


**Figure S3. HITS-Bio process.** Steps involved in loading spheroids. Spheroids were picked up from a spheroid chamber via selectively aspirating them and then transferred to a location for bioprinting onto a gel substrate. Red arrows demonstrate loaded spheroids.

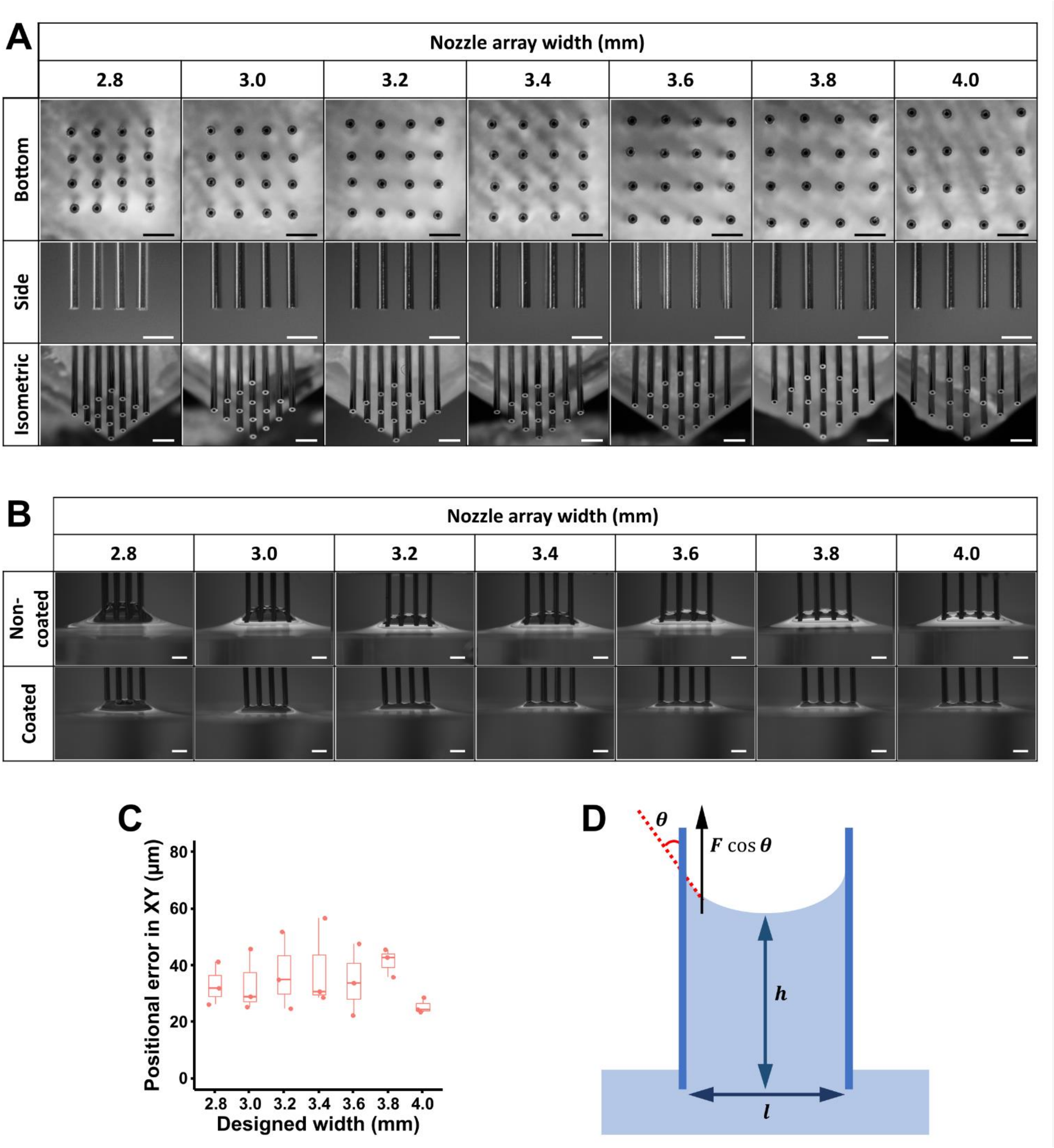


**Figure S4.** (A) Bottom, side, and isometric views of DCNA with varying designed widths. Scale bar: 1 mm. (B) Side view of liquid elevation in DCNA to evaluate the effect of silicon coating on DCNA. Scale bar: 1 mm. (C) Positional error in XY of DCNA with different designed widths (*n* = 3). (D) A schematic demonstration to describe the liquid elevation in DCNA and related parameters.

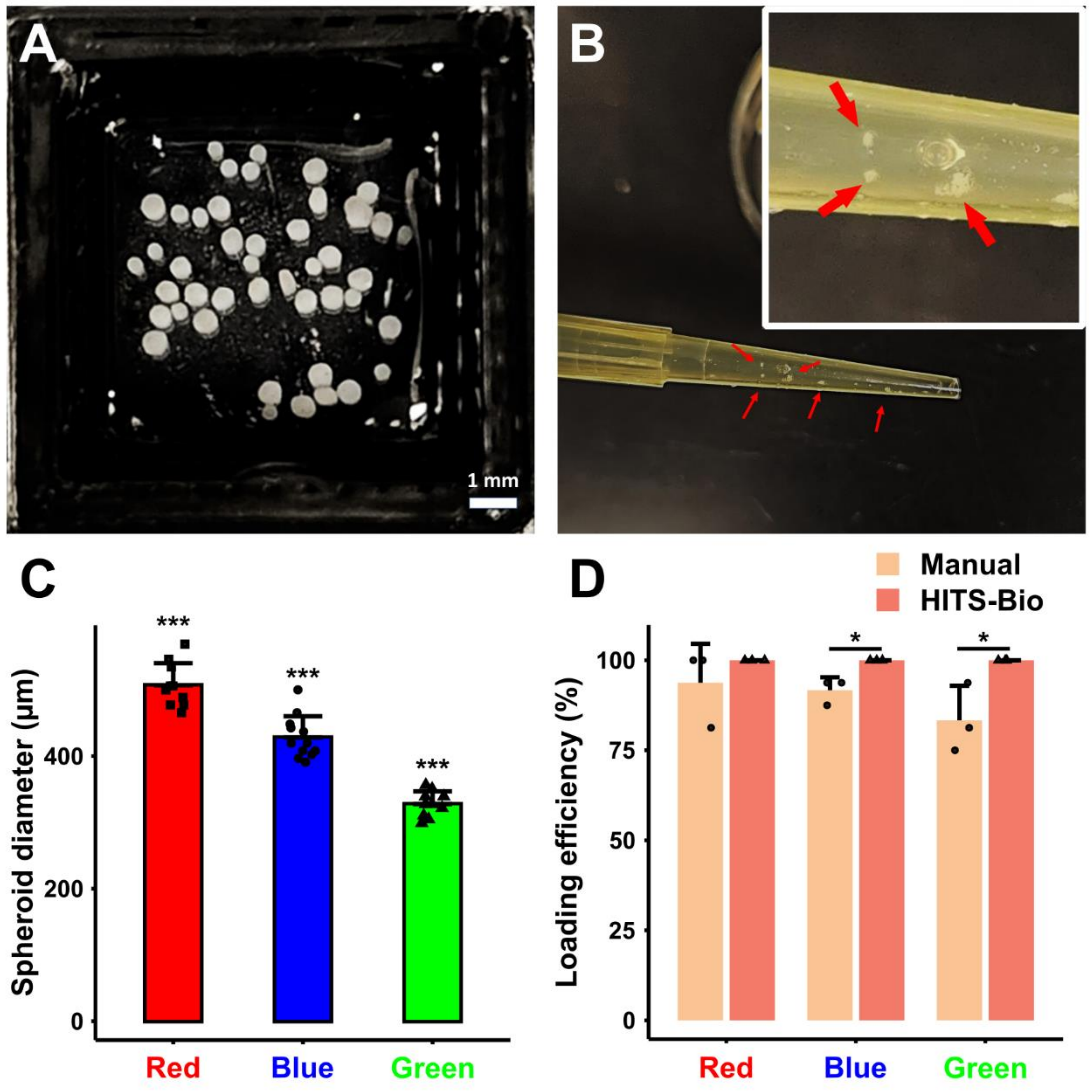


**Figure S5.** (A) Manually loaded different types (size and fluorescently labelled (color)) of spheroids loaded on 10% GM, followed by photo-crosslinking. (B) Red arrow indicates spheroids attached inside the pipette tip while mixing process without extrusion. (C) Diameter of spheroids stained with different colors (*n* = 10; ***p<0.001), (D) Comparison of loading efficiency (%) for manually loaded spheroids vs. HITS-Bio bioprinted spheroids for each color (*n* = 3; *p<0.05).

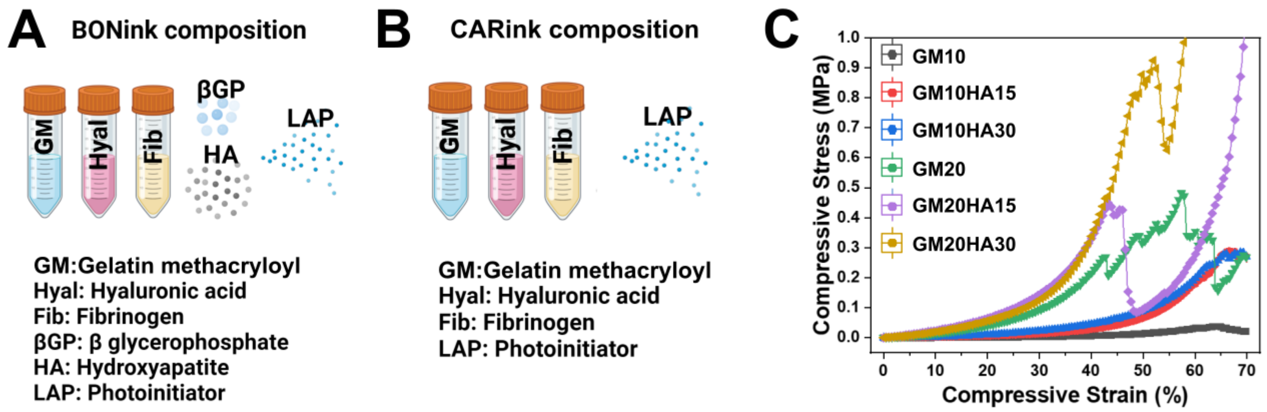


**Figure S6.** (A) BONink components, (B) CARink components, and (C) compressive stress of BONink and CARink with different concentrations of GM and HA.

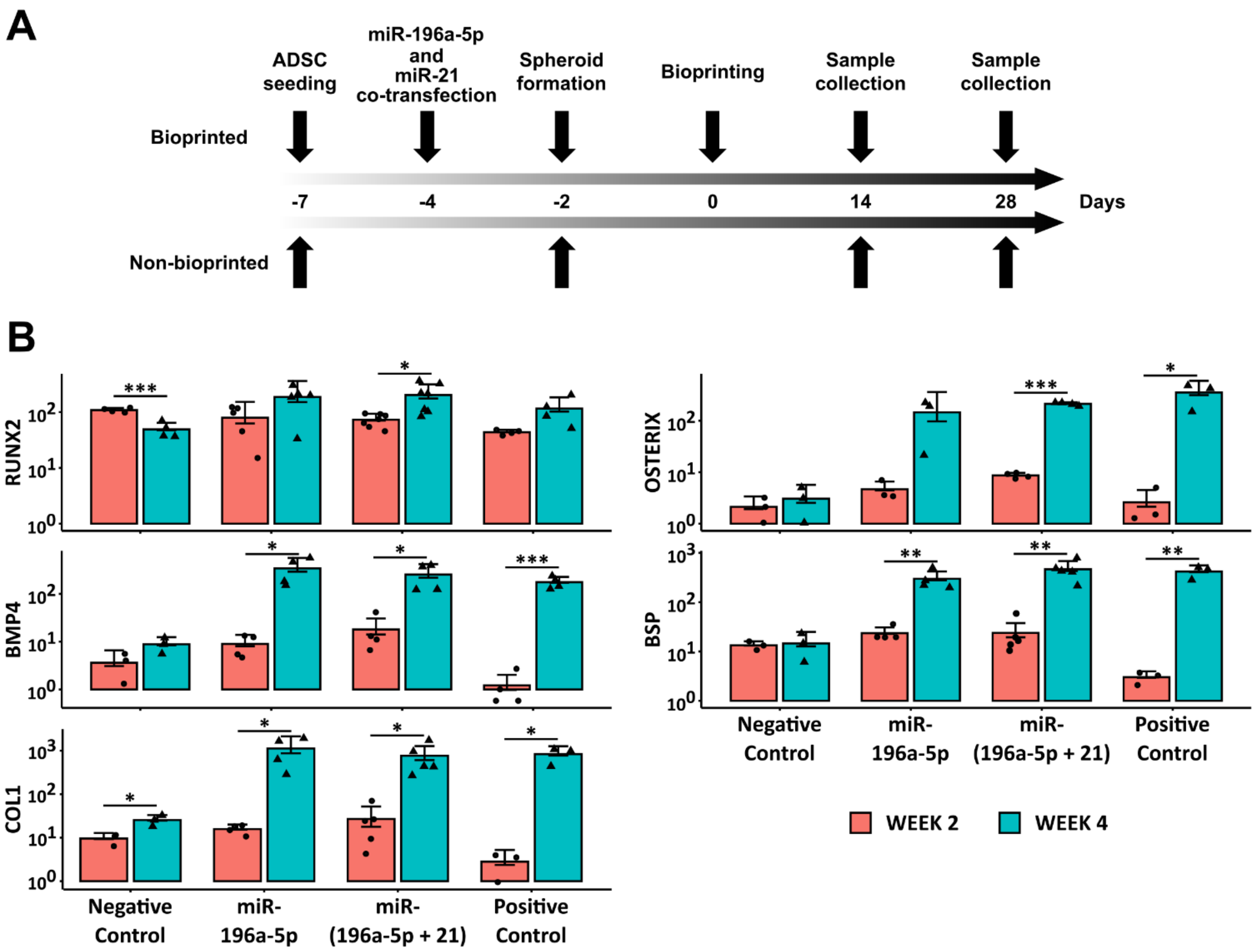


**Figure S7.** (A) A schematic describing the timeline and preparation of miRNA transfection and sample collection. (B) Quantitative gene expression of non-transfected ADSCs spheroids (negative control), miR-transfected spheroids and spheroids of ADSCs cultured with the osteogenic differentiation medium (positive control) at Weeks 2 and 4, normalized with respect to non-transfected ADSCs at Day 1 ($n$ = 4, *$p < 0.05$, **$p < 0.01$ and ***$p < 0.001$).

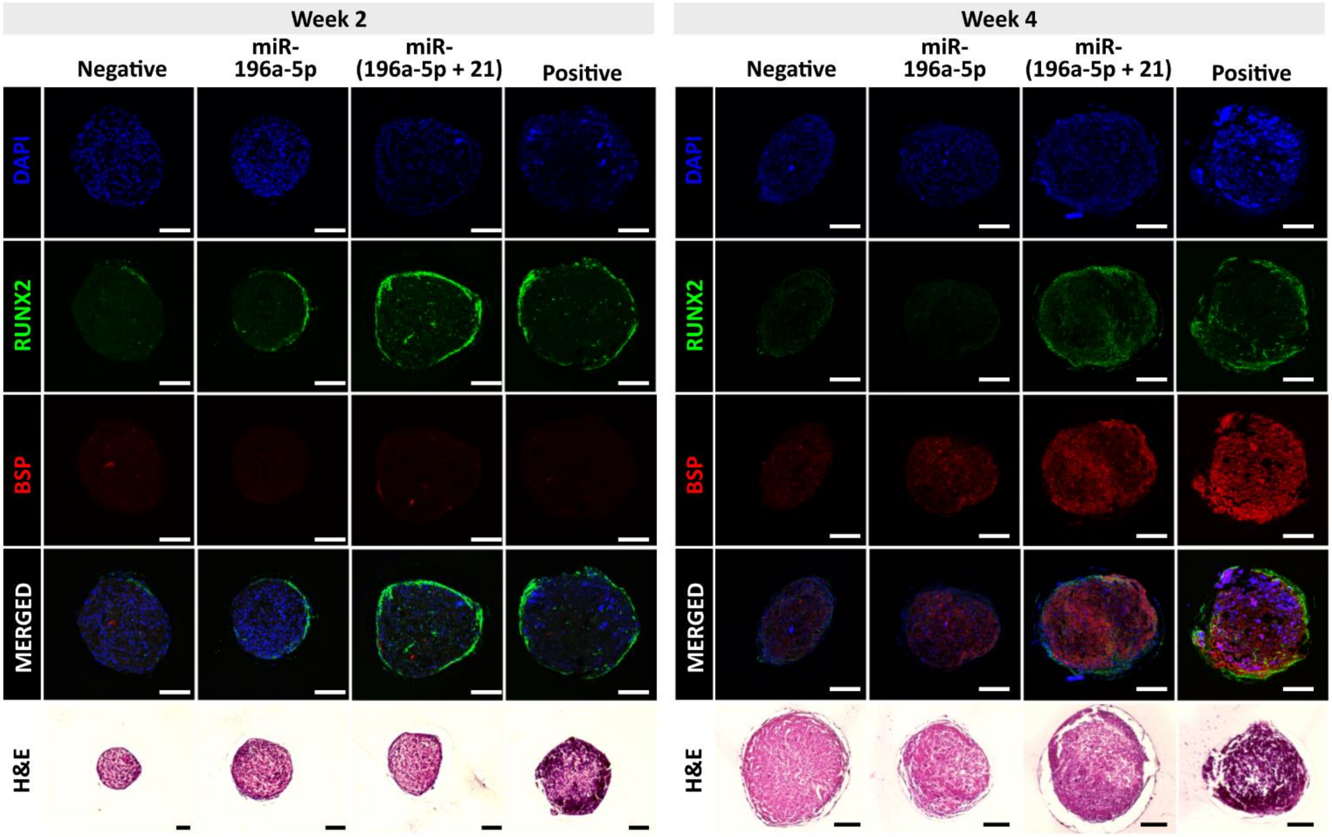


**Figure S8.** IHC images of non-transfected and transfected hADSC spheroids stained with RUNX2, BSP, and H&E at Weeks 2 and 4 (Scale bars, 200 µm).

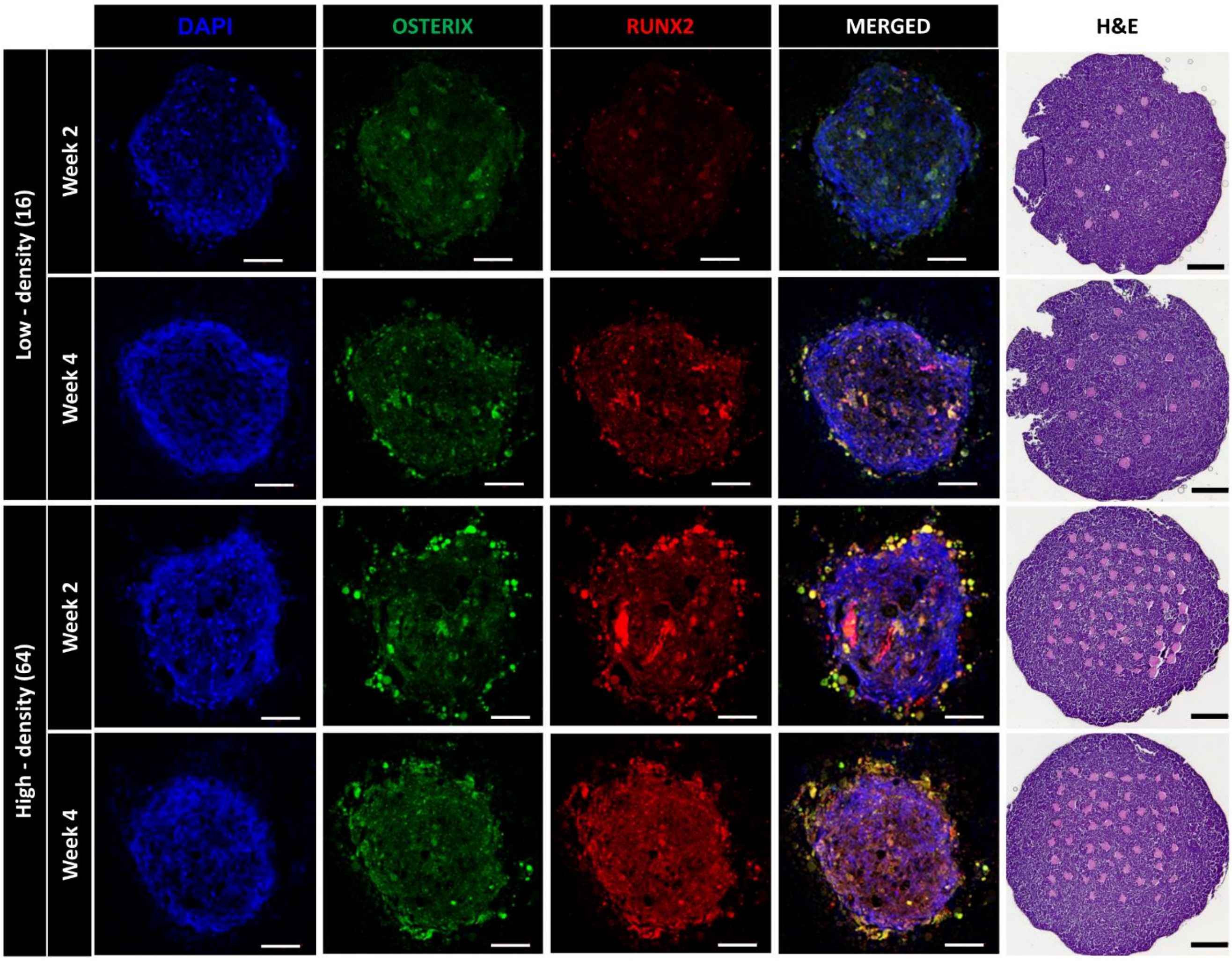


**Figure S9.** IHC (Scale bar: 100 µm) and H&E (Scale bar: 1 mm) images of bioprinted bone tissue with two different densities (low and high) at Weeks 2 and 4.

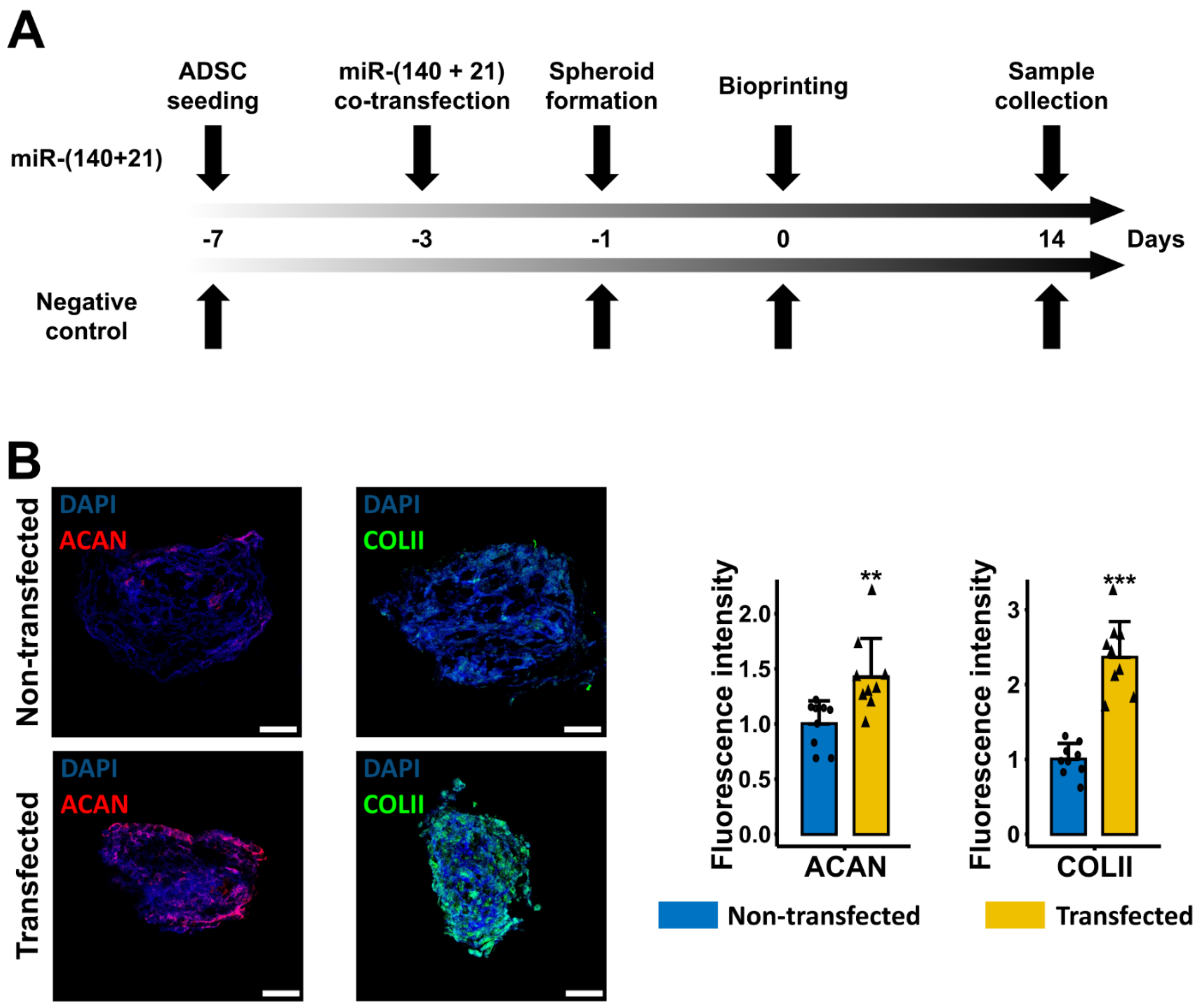


**Figure S10.** (A) A schematic describing the timeline and preparation of miRNA transfection and sample collection for SCT fabrication. (B) IHC with ACAN and COLII for non-transfected and transfected spheroids in $cm^3$ cartilage tissues at Week 2 and their fluorescence intensity ($n$ = 9, **$p$ < 0.01 and ***$p$ < 0.001). Scale bar: 50 μm.

**Table S1.** Fold change in gene expression for osteogenic marker on Week 4 as compared to Week 2, normalized to negative control (non-transfected hADSCs on Day 1).

| **Markers** | **2D Transfection** | **Transfected Spheroids** | **HITS-Bio Low Density** | **HITS-Bio High Density** |
|---|---|---|---|---|
| RUNX2 | 2.3 | 2.7 | 4.8 | 1.1 |
| BMP4 | 7.4 | 23.3 | 1.8 | 2.4 |
| COL1 | 2.3 | 52.1 | 5.1 | 2.5 |
| OSTERIX | 26.5 | 21.3 | 6.1 | 15.4 |
| BSP | 1.3 | 14.1 | 8.6 | 9.6 |

**Table S2.** Primers of the genes used in the qRT-PCR study.

| Gene | Forward primer | Reverse primer |
|---|---|---|
| COL1 | ATG ACT ATG AGT ATG GGG AAG CA | TGG GTC CCT CTG TTA CAC TTT |
| RUNX2 | GGT TAA TCT CCG CAG GTC ACT | CAC TGT GCT GAA GAG GCT GTT |
| BSP | AAC GAA GAA AGC GAA GCA GAA | TCT GCC TCT GTG CTG TTG GT |
| BMP4 | TAG CAA GAG TGC CGT CAT TCC | GCG CTC AGG ATA CTC AAG ACC |
| OSTERIX | CCT CTG CGG GAC TCA ACA AC | AGC CCA TTA GTG CTT GTA AAG G |
| GAPDH | CAC ATG GCC TCC AAG GAG TA | GTA CAT GAC AAG GTG CGG CT |

### Supplementary Information

#### S1. Capillary reaction in DCNA

A capillary reaction was observed between nozzles (**Figure S4D**), where liquid rises in a narrow gap against gravity. The surface tension of the liquid was calculated based on the height of elevated liquid, gravity, density of the liquid, and the inter-nozzle distance of DCNA. At equilibrium, the upward force must balance the downward force. The upward force pulls a specific volume of liquid upward, equivalent to the weight of the lifted liquid:

$$F_{st}\cos\theta = \gamma_{st}L = F_{weight} \quad (1)$$

where, $F_{weight}$ is force from the weight of the liquid pulled up, and $\gamma_{st}$ is the coefficient of surface tension of liquid (water- 0.0727 N/m at 20 °C). $L$ is the length of the boundary between the liquid and surface along with the surface tension acts (**Figure S4D**). This can be simplified as $F_{weight}$ is equal to $mg$, where $m$ is mass and $g$ is gravitational force. The liquid will continue to elevate until the force from the surface tension equals the weight of the liquid pulled up above the original liquid level:

$$\gamma_{st} \times 4l = mg \quad (2)$$

And since $m = \rho V$, where $\rho$ is the density and $V$ is the volume of the lifted liquid, equation (2) can be rewritten as:

$$\gamma_{st} \times 4l = \rho V g \quad (3)$$

Since $V$ is the volume surrounded by nozzles in DCNA and is equal to $l^2 h$, where $l$ is inter-nozzle distance and $h$ is the height of the elevated liquid:

$$\gamma_{st} \times 4l = \rho l^2 h g \quad (4)$$

Equation (4) can be rearranged to:

$$h = \frac{4\gamma_{st}}{\rho l g} \quad (5)$$

Equation (5) is the relationship between the surface tension, elevated liquid height, and inter-nozzle distance in DCNA. Here, two parameters can be altered to decrease *h*: either increasing $l$ or decreasing $\gamma_{st}$.

**Supplementary Videos**

**Supplementary Video 1.** HITS-Bio software operation.

**Supplementary Video 2.** DCNA with non-coated vs. coated nozzles.

**Supplementary Video 3.** DCNA with intra-nozzle distance of 2.8 mm vs. 3.4 mm.

**Supplementary Video 4.** Single nozzle AAB vs. HITS-Bio for bioprinting of 64 spheroids.

**Supplementary Video 5.** Patterning of spheroids. Selectively patterned spheroids stained with DAPI (blue), F-Actin (red), and F-Actin (green) using the DCNA platform with various configurations.

**Supplementary Video 6.** HITS-Bio for bone tissue bioprinting. Video demonstrates bioprinted osteogenic spheroids with low (16 spheroids) and high densities (64 spheroids) using BONink.

**Supplementary Video 7.** HITS-Bio using CARink. Video demonstrates bioprinted spheroids with low (16 spheroids) and high densities (64 spheroids) on a transparent gel (CARTink) for clear visualization.

**Supplementary Video 8.** Intraoperative bioprinting (IOB) of bone tissue into critical-sized rat calvarial defects with BONink only, low-density (16 spheroids) and high-density (64 spheroids) groups. The process included EBB of the BONink, then deposition of miR-(196a-5p + 21) co-transfected osteogenic spheroids followed by another layer of BONink.

**Supplementary Video 9.** HITS-Bio for fabrication of 1 $cm^3$ cartilage constructs. The process took 40 min in total including the EBB of the CARink and HITS-Bio of miR-(140 + 21) co-transfected chondrogenic spheroids.
